# Spatial patterning of contractility by a mechanogen gradient underlies *Drosophila* gastrulation

**DOI:** 10.1101/2025.04.11.648359

**Authors:** Gayatri Mundhe, Valentin Dunsing-Eichenauer, Jean-Marc Philippe, Élise Da Silva, Claudio Collinet, Thomas Lecuit

**Affiliations:** Aix Marseille Université, CNRS, IBDM UMR7288, Turing Center for Living Systems, Marseille, France; Collège de France, Paris, France

## Abstract

During development cell deformations are spatially organized, however, how cellular mechanics is spatially controlled is unclear. Spatial control of cell identity often determines local cellular mechanics in a two-tiered mechanism (*1*). Theoretical studies also proposed that molecular gradients, so called “mechanogens”, spatially control mechanics (*2, 3*). We report evidence of such a “mechanogen” required for *Drosophila* gastrulation. We show that the GPCR ligand Fog, expressed in the posterior endoderm (*4–6*), diffuses and acts in a concentration-dependent manner to activate actomyosin contractility at a distance during a wave of tissue invagination. While Fog is uniformly distributed in the extracellular space, it forms a surface-bound gradient that activates Myosin-II via receptor oligomerization. This activity gradient self-renews as the wave propagates and is shaped by both receptor endocytosis and a feedback mechanism involving adhesion to the vitelline membrane by integrins. This exemplifies how chemical, mechanical and geometrical cues underly the emergence of a self-organized mechanogen activity gradient.

## Introduction

Embryonic development entails the spatial and temporal control of both cell fate determination to generate different cell types (*7*) and of cell mechanics to generate tissue shapes during morphogenesis (*1, 8, 9*). Classically, the latter is considered to be a consequence of the former. For instance, the vertebrate nervous system is first specified in the dorsal region of the ectoderm and subsequently shaped into the neural tube. Similarly, the *Drosophila* mesoderm is first specified in the ventral part of embryos, and then invaginates. Typically, spatial patterning in a field of cells involves morphogens, i.e. diffusing molecules (*10*) that form concentration gradients (*11*) and impart positional information within tissues (*12*). Morphogens are produced from a source from which they diffuse, are degraded within the target tissue and thereby produce exponential gradients (*13, 14*). Receiving cells interpret the positional information imparted by morphogen gradients by activating the expression of different genes at different morphogen concentration thresholds. This mechanism partitions the field of receiving cells in discrete domains with different cell identities (*15, 16*). For example, the morphogens Shh, BMP and Wnt specify the identity of different neuron progenitors in the vertebrate neural tube through the expression of different combinations of transcription factors (*17*). In *Drosophila*, the morphogen Dorsal specifies mesoderm identity on the ventral side of embryos via the expression of the transcription factors Twist and Snail (*18, 19*). Within this framework, the specification of cell identities directs downstream morphogenetic processes through the regulation of cell mechanics, such as, cell contractility, cell motility and cell adhesion (*1, 20*). For example, the differential expression of adhesion molecules downstream of the Shh morphogen controls sorting of neural progenitors in the zebrafish neural tube (*21*). In the *Drosophila* ventral mesoderm, the transcription factors Twist and Snail, in response to the morphogen Dorsal, induce the expression of Fog, a GPCR ligand that leads to the apical activation of the small GTPase Rho1, the kinase Rok and non-muscle Myosin II (MyoII) (*4, 22–25*). This induces cell autonomous apical constriction and culminates with the invagination of the mesoderm in the ventral part of embryos. Thus, morphogens direct morphogenesis via a two a two-tiered mechanism whereby the spatial control of cell identities determines downstream local cellular mechanics.

By analogy to the concept of morphogens, theoretical studies (*2, 3*) proposed that so-called “mechanogens”, i.e. “external diffusible biomolecules”, may form spatial gradients that “influence mechanical properties, such as cell-cell adhesion and cellular contractility (and therefore, cell shape), and create spatial gradients in cell structure in a tissue” (*3*). Recent studies reported that classical morphogens can also function as mechanogens by regulating collective tissue mechanical properties without specifying cell fates. For instance, a gradient of Nodal in Zebrafish specifies mesendoderm invagination by 1. triggering a motility-driven unjamming transition in protrusive leader cells able to autonomously internalize, and 2. promoting adhesion to their immediate followers, which results in a collective and ordered mode of internalization (*26*). Similarly, in the developing feather buds, the morphogens BMP and FGF define two adjacent domains with different supracellular mechanical properties, a rigid elastic core and an active fluid-like margin, that drive mesenchymal budding (*27*). In both cases, the morphogens Nodal and BMP can be considered to act as mechanogens as they tune collective, supracellular mechanical properties, however, they most likely do it indirectly via transcriptional regulation of target genes. Instead, mechanogens have been proposed to form chemical gradients that directly regulate cell mechanics (e.g. cell contractility) without the intermediary of gene transcription (*2*). Here, we report evidence that Fog, a signaling molecule directly controlling cellular contractility, in addition to its autocrine function in the *Drosophila* posterior midgut anlage, acts in a concentration-dependent manner to control MyoII activation at a distance during the propagation of a wave of tissue invagination. As such, we propose that Fog functions as a bonafide concentration-dependent mechanogen. We also study the mechanisms of mechanogen gradient formation.

## Results

### GPCR signalling is required in the propagation zone for MyoII activation

The morphogenesis of the *Drosophila* posterior endoderm begins with MyoII-dependent apical constriction in the posterior-most region of the embryo, the endoderm primordium. Here the terminal transcription factors Hückebein and Tailless (*28*) control the localized expression and secretion of the ligand Fog which activates tissue autonomous recruitment of MyoII apically and tissue bending (*4–6*). This induces a polarized flow towards the embryo dorsal-anterior due to the high curvature of the tissue in this region (*29*) (Fig. 1A,B). This initial flow and tissue bending triggers a wave of MyoII activation and tissue invagination that propagates anteriorly within the dorsal epithelium (Fig.1A and 1C). Wave propagation does not depend on sustained gene transcription and is driven by a self-replicating cycle of 3D cell deformation (Fig. 1C) where posterior tissue invagination induces in a sequence: 1) contact with the vitelline membrane (VM), 2) integrin-mediated adhesion, 3) MyoII activation and 4) detachment from the vitelline membrane of the more anterior cells in the wave propagation zone (PZ) (*6*) due to contractility in the furrow (*30*). During the wave, MyoII is initially recruited at low levels in cells distant from the invaginating furrow upon their contact with the vitelline membrane. Then, when cells get closer to the furrow, MyoII recruitment increases in a positive feedback loop with integrins (*30*). Although previous studies (*6*) ruled out that the propagation of MyoII activation reflects the propagation of a wavefront of diffusion/transport of the ligand Fog, the role of Fog-GPCR signalling in activating MyoII in cells of the PZ was not directly tested. Furthermore, the absence of integrins (KO of the *⍺*-PS3 integrin, Scab) does not completely abolish MyoII activation, which remains pulsatile at low levels, during wave propagation (*30*), indicating that other factors, such as Fog-GPCR signalling, could be involved.

**Figure 1.**
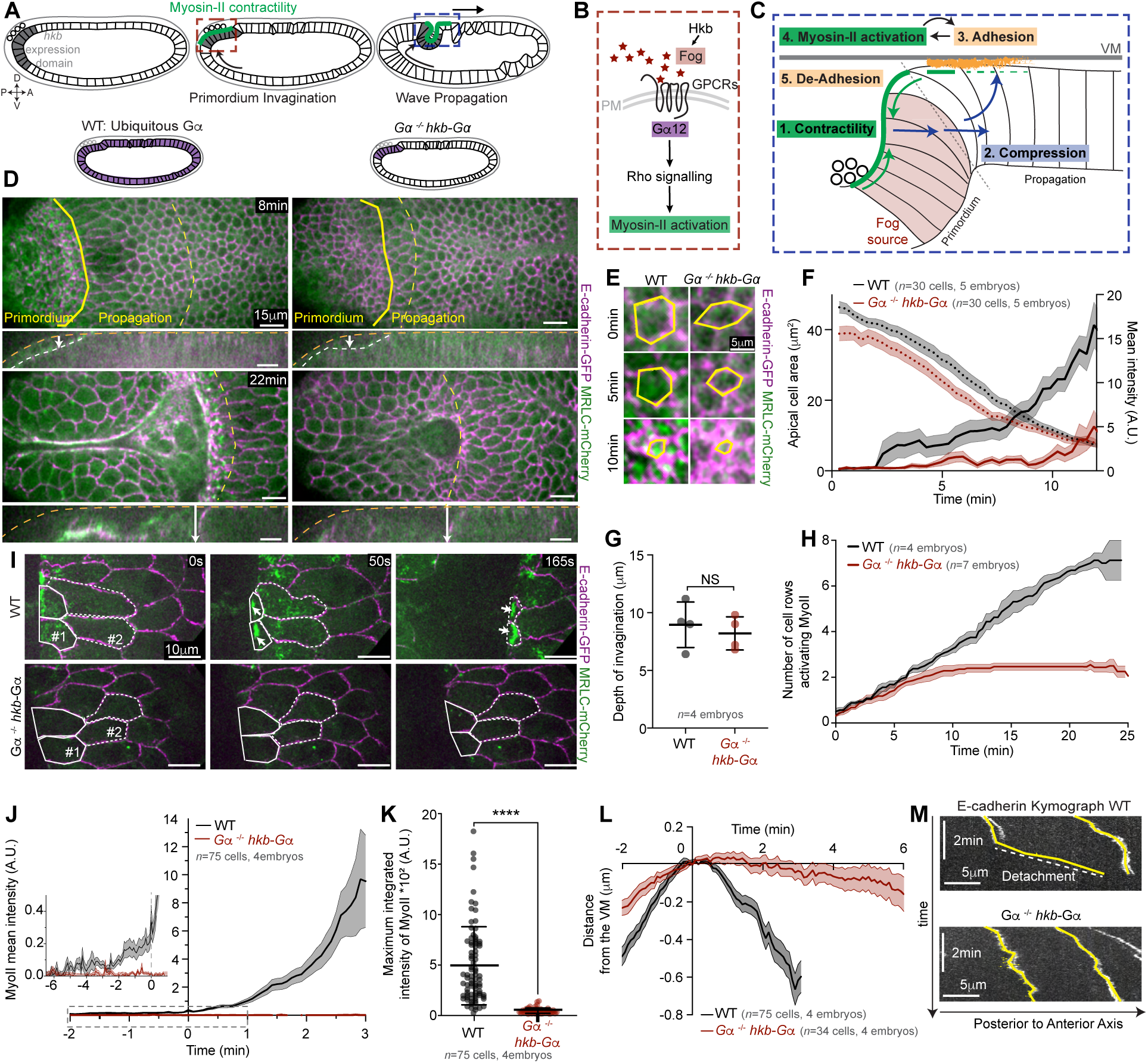
MyoII activation in the PZ requires GPCR signalling. (**A**): Cartoon of endoderm invagination during *Drosophila* gastrulation. The grey cells indicate the domain of *hückebein* (*hkb*) expression and the green line represents the region of MyoII contractility. (**B**): Activation of MyoII by Fog/GPCR signalling during primordium invagination (dashed red box in 1A). (**C**): Schematic of the 3D cell deformation cycle during MyoII wave propagation (dashed blue box in 1A). MyoII contractility (green arrows) generates compressive forces (blue arrows) leading to integrin-dependent adhesion (orange zone) which activates MyoII. Feedback amplification between adhesion and MyoII activation (black arrows), VM: Vitelline membrane. (**D**): Top and side views from a time-lapse of the posterior endoderm invagination in a WT (*top*) and a *Gα*^−/−^ *hkb*-Gα (*bottom*) embryos, imaged with 40X objective. The yellow solid and dashed lines highlight the anterior limits of the primordium and propagation zone, respectively. In side views the yellow dotted line indicates the vitelline membrane. White arrows indicate the depth of the invagination. Time 0 is defined as 22 min prior to first cell division (see methods). On the top, a drawing indicating the regions in the embryo where Gα is expressed (in purple). (**E**): Close-up views over time of representative cells in the primordium in the indicated conditions. In yellow the contours of a cell tracked over time. (**F**): Average time traces of apical cell area and MyoII mean intensity in cells of the primordium region. *n=*30 cells from 5 embryos each. (**G**): Measurements of the depth of the primordium invagination at T=8 mins in WT and *Gα*^−/−^ *hkb*-Gα embryos. *n=*4 embryos each. *P*=0.17, unpaired *t*-test with Welch’s correction. (**H**): Cumulated number of cell rows activating MyoII over time in WT and *Gα*^−/−^ *hkb*-Gα embryos. Time 0 corresponds to the onset of MyoII activation in the first cell of the PZ for each embryo. *n=*4 embryos for control and 7 for *Gα*^−/−^ *hkb*-Gα. (**I**): Stills from a high resolution (100X objective) time-lapse of MyoII activation in the PZ. Representative cells are tracked over time (white solid and dashed line). White arrows indicate activation of MyoII at high levels before cell detachment. (**J**): MyoII mean intensity over time in cells of the PZ in the indicated conditions. Time 0 for each cell is the time when the cell area is maximum (see methods). Inset: blow-up of the region in the dashed box. (**K**): Maximum integrated intensity of MyoII in cells of the PZ for the indicated conditions. \*\*\*\**P*<0.0001, unpaired *t*-test with Welch’s correction. (**L**): Temporal profile of cell detachment from the VM quantified as the average distance of E-cadherin junctions from the VM for each cell. *n=*75 cells from 4 embryos in both conditions in **J** and **K** while *n=*75 cells in WT and 34 in *Gα*^−/−^ *hkb*-Gα from 4 embryos each in **L**. (**M**): Kymographs along the AP-axis of E-cadherin junctions in representative cells in the indicated conditions. A, anterior; P, posterior; D, dorsal; V, ventral. Statistics: Data in **G** and **K** is mean±s.d., data in **F**, **H**, **J** and **L** are mean±s.e.m.. Scale bars, 15 μm (**D**), 10 μm (**I**) and 5 μm (**E**, **M**).

The small GTPase Gα12/13 Concertina (hereafter Gα) is known to transduce Fog-GPCR signalling that activates medio-apical MyoII contractility and tissue invagination in the endoderm and mesoderm primordia (*4, 25, 31, 32*). In the absence of Gα (null mutants *Gα^−/−^*), invagination of the endoderm primordium, which triggers the subsequent wave propagation, is blocked (*6*). Thus, to test the role of Fog-GPCR signalling in cells of the PZ, we rescued Gα activity in *Gα^−/−^* embryos only in the primordium to be able to induce its invagination. To this end, we expressed a WT form of Gα using a *hückebein* (*hkb*) promoter, which drives expression specifically in the posterior endoderm primordium and not in the neighbouring PZ (*32, 33*), in *Gα^−/−^* embryos (*Gα^−/−^ hkb*-Gα hereafter). We visualized cell contours with E-cadherin::GFP and MyoII activation with MyoII Regulatory Light Chain MRLC::mCherry. As expected, in *Gα^−/−^ hkb-Gα* embryos, cells in the primordium activated apical MyoII and constricted apically (Fig. 1D-F, Fig. S1A and Movies S1 and S2). This led to the formation of an initial invagination with kinetic similar to those in WT embryos (Fig. 1D side views and 1G), indicating a functional rescue of Gα activity in the primordium. However, despite a normal invagination of the endoderm primordium, in Gα^−/−^ *hkb*-Gα embryos the subsequent wave propagation (Fig. 1H-L) and the dorsal-anterior movement of the posterior endoderm (Fig. S1B,C and Movie S3) were severely affected. Indeed, MyoII activation occurred on average only in 2.5±0.06 s.e.m. rows of cells compared to an average of 7.1±0.3 s.e.m. cells in WT embryos (Fig. 1H). Beyond the first two cells at the border with the endoderm primordium, in the PZ of *Gα^−/−^ hkb*-Gα, MyoII was not activated at all (Fig.1I-K and Movie S4). As a result, non-contractile cells joined the furrow, which blocked detachment from the VM of more anterior cells once they reached the edge of the invaginating furrow (Fig. 1I and 1L-M). Together, we conclude that Gα signalling in cells of the PZ is required to activate MyoII and sustain wave propagation.

### Fog diffusion from the primordium is necessary for MyoII activation in the propagation zone

In *Drosophila*, Gα signalling inducing MyoII-dependent tissue invagination is activated by the secreted GPCR-ligand Fog (*4, 5*). We thus considered that Fog might be the ligand activating Gα signalling in the PZ. Consistent with this possibility, Fog protein was detected by immunohistochemistry in the PZ (*6*). In the posterior endoderm, *fog* is expressed zygotically at high levels only in the primordium region (*4, 6*) but maternally deposited *fog* transcripts have been detected at low levels everywhere in the embryo (*4*) and could give rise to Fog detected in the PZ. To test the roles of these two possible sources of Fog, we generated embryos where Fog is expressed only zygotically in the endoderm primordium. To this end, we blocked both maternal and zygotic expression of the endogenous gene by injection of dsRNAs against *fog* (*fog* RNAi) (*6, 34*) and restored its expression only in the endoderm primordium using an RNAi-resistant *fog* transgene expressed under the *hkb* promoter (hereafter *hkb-fog_res_*) (Fig. 2A and Fig. S2A). Contrary to the expression of an RNAi-sensitive *hkb-fog* transgene (*hkb-fog_sen_* hereafter), expression of *hkb*-*fog_res_* in *fog* RNAi embryos rescued the invagination and the dorsal-anterior movement of the posterior endoderm (Fig. 2D top and middle, Fig S2B-E and Movies S5 and S6), indicating that *fog_res_* is indeed resistant to RNAi. Notably, *hkb*-*fog_res_* rescued apical MyoII recruitment, cell apical constriction and tissue invagination, which were defective in *hkb-fog_sen_* embryos in the endoderm primordium (Fig. 2D-G and Movie S7). Furthermore, *hkb*-*fog_res_* also rescued the propagation of MyoII activation in *fog* RNAi embryos. Propagation of MyoII activation in these embryos spanned on average 5.3±0.2 s.e.m. cells (Fig. 2H) while it was completely blocked in *hkb-fog_sen_* embryos injected with *fog* dsRNAs (Fig. 2D). Moreover, in *hkb*-*fog_res_* embryos injected with *fog* dsRNAs the levels of MyoII were almost as high as in *hkb*-*fog_res_* embryos injected with water (Fig. 2I,J and Movie S8), showing that *fog* expression in the endoderm primordium is sufficient to rescue endoderm invagination and wave propagation in *fog* RNAi embryos. We conclude that Fog produced in the endoderm primordium acts non-autonomously in the PZ to activate GPCR-dependent MyoII activation there.

**Figure 2.**
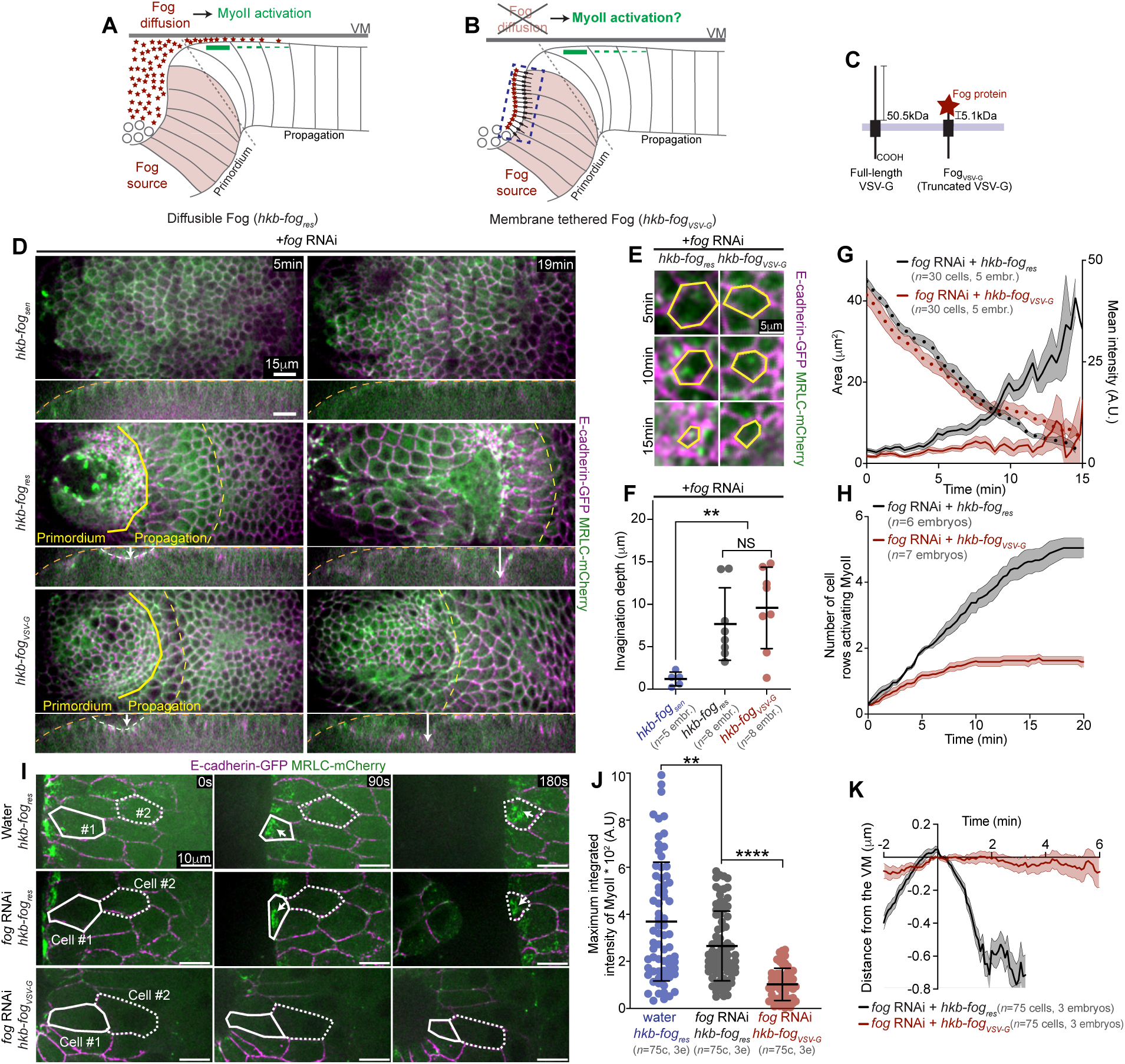
Fog production from the primordium and its dispersion into the PZ are necessary for MyoII activation. (**A-B**): Schematics of the expression of an RNAi-resistant diffusible (**A**) and membrane-tethered version (**B**) of Fog under the *hkb* promoter in a cross-section of the dorsal epithelium. The red stars indicate Fog, and red shaded cells indicate the endoderm primordium where the *hkb* promoter is active. (**C**): Schematic of the membrane-tethered Fog fused with a truncated form of type-II transmembrane protein VSV-G used in this study. The full-length version of VSV-G is also shown on the left as a comparison. (**D**): Top and side views from time-lapses of the posterior endoderm invagination in embryos injected with *fog* RNAi and expressing under the *hkb* promoter an RNAi-sensitive (*hkb-fog_sen_*, top), an RNAi-resistant (*hkb-fog_res_*, middle) or an RNAi-resistant version of *fog* tethered to the plasma membrane (*hkb-fog_VSV-G_*, bottom). The yellow solid and dashed lines highlight the anterior limits of the primordium and propagation zone, respectively. In side views, the white arrows indicate the depth of the invagination. A yellow dotted line indicates the vitelline membrane. (**E**): Close-up views over time of representative cells in the primordium in the indicated conditions. In yellow the contours of a cell tracked over time. (**F**): Quantification of the primordium invagination depth at T=5 min in the indicated conditions. *N=* 5 embryos for *hkb-fog_sen_* and 8 embryos for both *hkb-fog_res_* and *hkb-fog_VSV-G_.* \*\**P*=0.0033 (*hkb-fog_sen_ vs hkb-fog_res_*), \*\**P*=0.0015 (*hkb-fog_sen_ vs hkb-fog_VSV-G_*) and NS, *P*=0.4159 (*hkb-fog_res_ vs hkb-fog_VSV-G_*) with unpaired *t*-test with Welch’s correction. (**G**): Average time traces of cell apical area and MyoII mean intensity in primordium cells. *N=*30 cells from 5 embryos each. (**H**): Cumulated number of cell rows activating MyoII in the PZ over time in *hkb-fog_res_* and *hkb-fog_VSV-G_* embryos injected with *fog* RNAi. *N=*6 and 7 embryos in *hkb-fog_res_* and *hkb-fog_VSV-_*_G_, respectively. (**I**): Stills from high-resolution time-lapses of MyoII activation in the PZ in *hkb-fog_res_* and *hkb-fog_VSV-G_* embryos injected with water (control) or *fog* RNAi. Representative cells are outlined in white and tracked over time. White arrows indicate activation of MyoII at high levels before cell detachment. (**J**): Maximum MyoII integrated intensity in cells of the PZ in the indicated conditions. \*\**P*=0.0056 and \*\*\*\**P*<0.0001, unpaired *t*-test with Welch’s correction. (**K**): Temporal profile of detachment from the VM quantified as the distance of E-cadherin junctions from the VM. *n=*75 cells from 3 embryos each in **J** and **K**. Statistics: Data in **F** and **J** are mean±s.d. and data in **G**, **H** and **K** are mean±s.e.m. Scale bars, 15 μm (**D**), 10 μm (**I**) and 5 μm (**E**).

Next, we investigated the mechanism by which Fog acts at a distance in the PZ. Fog is a large (100kDa), secreted molecule and previous reports did not test whether Fog acts in a paracrine manner (*4, 5*). In light of the non-autonomous rescue of MyoII activation in the PZ reported above, we hypothesized that Fog produced and secreted in the primordium might diffuse into the PZ to activate MyoII. To test this, we sought to inhibit Fog dispersal from its source of production (Fig. 2B). We constructed a membrane-tethered version of Fog by fusing the entire *fog* coding sequence with a truncated form (Δ1-421, see methods for details) of the Vesicular Stomatitis Virus (VSV) G protein, a type-II transmembrane protein (Fig. 2C), and tested its ability to disperse from a localized production source. We expressed Fog-VSV-GΔ1-421 (Fog_VSV-G_ hereafter) in stripes in the embryo using the *wingless-Gal4* driver (Fig. S3A) and visualized it with an anti-Fog antibody (*35*). Contrary to a WT version of Fog that was detected also into the neighbouring non-expressing tissue, Fog_VSV-G_ was only observed within the producing cells (Fig. S3B,C). To test the ability of Fog_VSV-G_ to activate MyoII, we overexpressed it homogeneously in the embryo during gastrulation. The resulting pattern of MyoII activation, with low-level activation in the entire dorsal epithelium and strong MyoII activation in the endoderm primordium and the PZ (*6*), was similar from that of an overexpressed WT Fog (Fig. S3D,E and Movie S9). This confirmed that Fog_VSV-G_, although unable to diffuse, is able to activate GPCR signalling and MyoII contractility cell-autonomously in the embryo. We next used this tool to test whether Fog produced in the primordium diffuses to the PZ to activate MyoII there. We expressed an RNAi-resistant version of Fog_VSV-G_, in the primordium (using the *hkb* promoter) in embryos injected with *fog* dsRNA to remove endogenous maternal and zygotic Fog (hereafter referred to as *hkb-fog_VSV-G_*, Fig. 2B). In *hkb-fog_VSV-G_* embryos, cell apical constriction and endoderm primordium invagination were similar to *hkb-fog_res_* (Fig. 2D-G and Movies S5 and S7). However, the subsequent anterior movement of the endoderm was blocked (Fig. S2F,G and Movie S6) as was wave propagation (Fig. 2D,H, on average only 1.7±0.1 s.e.m cells). Indeed, in *hkb-fog_VSV-G_* embryos MyoII was not activated in cells of the PZ beyond the first 1-2 cell rows (Fig. 2I,J and Movie S8), and cells did not detach from the VM (Fig. 2K).

Altogether, we conclude that Fog produced in the primordium diffuses into the zone of wave propagation where it is required to activate MyoII and cell detachment. Our findings demonstrate that Fog acts in a tissue non-autonomous manner during wave propagation.

### Fog production and uptake tune an exponential gradient of MyoII activity

Since Fog diffuses from the primordium to activate MyoII at a distance, we next explored the spatial dynamics of Fog activity. During wave propagation, apical MyoII is graded in a restricted domain of 2-3 cell rows over ∼30-40 µm from the advancing furrow, with high MyoII levels in cells just anterior to the furrow and lower levels more anteriorly (*30*). We considered the possibility that Fog diffusion from the primordium and degradation or removal in the receptive field may set the amplitude and range of the graded activation of MyoII similar to morphogen activity gradients (*14, 36*). We found that at each moment in time, as the invaginating furrow moves towards the anterior, the spatial distribution of MyoII mean intensity in front of the furrow fits a one-phase exponential decay curve with an average length-scale (λ) of 3.18±0.29 s.e.m μm. (Fig. 3A-C, see methods). The shape of the activity gradient of morphogens typically depends on 1) the rate of morphogen production at the source, 2) the rate of its removal in the target tissue and 3) its diffusion coefficient. We tested whether similar mechanisms also control the gradient of MyoII activity in the PZ. To this end, we first increased Fog production in the primordium (the source) by expressing two additional copies of a WT form of Fog using the *hkb* promoter. This increased both the amplitude in front of the invaginating furrow (at x=0 µm, WT: 11.55±0.10 A.U and *hkb-fog* 32.59±0.32 A.U, mean±s.e.m.) and the range of the gradient (WT: 27.08±5.72 µm, *hkb-fog:* greater than 38.01±1.92 µm, mean±s.e.m.) as expected of a concentration gradient that defines the activation profile of MyoII (Fig. 3D-F and Supplementary Video 10). Interestingly, the length scale of the exponential MyoII activity gradient also increased (WT: 3.18±0.29 μm and *hkb-fog*: 5.26±0.58 μm, mean±s.e.m.), suggesting that diffusion or degradation/removal of Fog may depend on Fog concentration. Next, we tested whether the spatial activity of MyoII also depends on Fog removal within the PZ. Since Fog binding to its GPCR induces endocytosis of the receptor-ligand complex (*37*), we sought to interfere with Fog removal by blocking GPCR endocytosis. Gprk2, a kinase that promotes GPCR endocytosis upon ligand binding (*38, 39*), is the only GRK known to interfere with Fog signalling and endocytosis in the embryo (*35, 37*). Knock-down of Gprk2 by RNAi (*gprk2* RNAi) resulted in an increase in the amplitude (Control: 9.72±0.08 A.U and *gprk2* RNAi: 14.25±0.14 A.U, mean±s.e.m.), the range (Control: 20.86±5,72 µm and gprk2 RNAi: greater than 33.59±2.5 µm, mean±s.e.m.) and length-scale (Control: 2.96±0.45μm and *gprk2* RNAi: 5.48±0.58μm, mean±s.e.m.) of the exponential MyoII activation profile (Fig. 3G-I and Movie S11) similar to *hkb-fog*, indicating that Fog removal/endocytosis also tunes the MyoII activity gradient.

**Figure 3.**
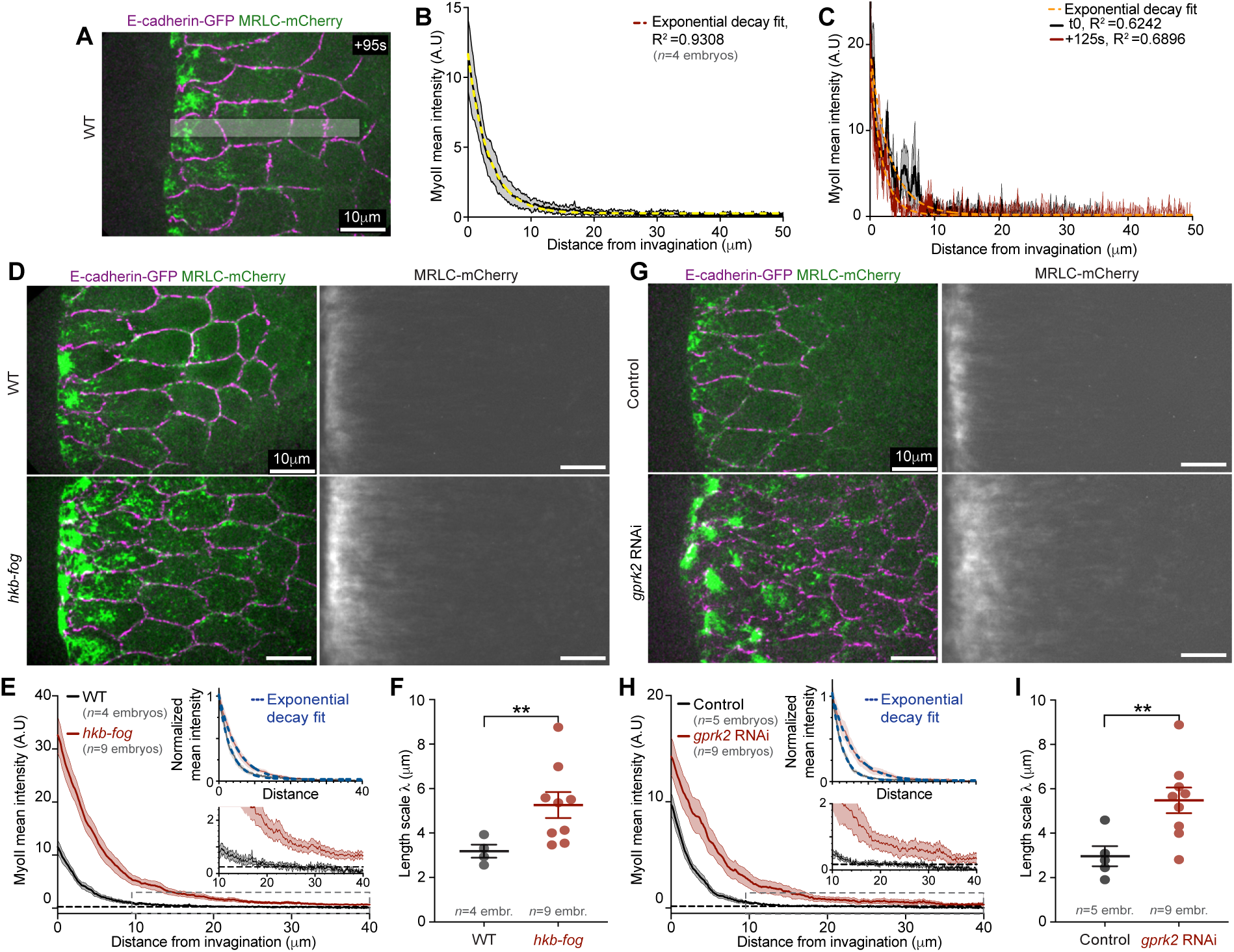
MyoII activation follows and exponential gradient tuned by Fog production in the primordium and removal in the PZ. (**A**): Still image from a high-resolution time-lapse of MyoII in the PZ in a WT embryo. (**B**): Average spatial profile of MyoII mean intensity in the PZ in WT embryos. *n*=4 embryos. The yellow dashed line is the fit with a single exponential decay function. (**C**): Spatial profile of MyoII mean intensity at different time points from a single embryo. The orange dashed lines are the fit with a single exponential decay curve. In **B** and **C** R^2^ is the coefficient of determination and measured the goodness of the fit to a one-phase exponential decay curve (see methods). (**D**): Left: Still images from high-resolution time lapses of MyoII in the PZ in WT (top) and embryos with increased Fog expression in the primordium (*hkb-fog*). Right: Temporal averaging of MyoII intensity in the referential of the moving invagination (see methods). (**E**): Average spatial profile of MyoII mean intensity in the PZ in the indicated conditions. The dashed horizontal line indicates the activation threshold for MyoII estimated as described in the methods. The inset at the bottom is a zoomed view of the dashed box, The inset at the top shows the same profiles normalized using min-max normalization (see methods). (**F**): Decay length scale of the MyoII intensity spatial profile in individual embryos of the indicated conditions. \*\**P*=0.0089, unpaired *t*-test with Welch’s correction. (**G-I**): Identical set of images and quantification as described in **D**-**F** for control (top) and embryos uniformly expressing shRNAs against Gprk2 (*gprk2* RNAi, bottom). \*\**P*=0.0051, unpaired *t*-test with Welch’s correction. **E** and **F,** *n*=4 embryos (WT) and 9 embryos (*hkb-fog*). **H** and **I,** *n*=5 embryos (Control) and 9 embryos (*gprk2* RNAi). Statistics: Data in **F** and **I** are mean±s.d. and data in **B**, **C**, **E** and **H** are mean±s.e.m. Scale bars: 10 μm.

Altogether, these data show that the Fog produced in the endoderm primordium sets an exponential activity gradient of MyoII in the PZ by diffusion/degradation and thus acts as a mechanogen directly controlling cellular contractility (*2, 3*). It is worth noting that this exponential gradient, whose maximum is located at the edge of the tissue invagination, is not static but translocates anteriorly as the wave of invagination propagates to the anterior.

### Fog is uniformly distributed in the extracellular space

Since Fog regulates the gradient of MyoII activation in the PZ, we next tested whether Fog also forms a concentration gradient there (Fig. 4A). Immunohistochemistry (IHC) against Fog in fixed embryos revealed a concentration gradient of cellular Fog protein in the PZ, although shallower than that of MyoII (Fig. 4B,C), consistent with the integrin-dependent amplification of MyoII activation closer to the invaginating furrow (*30*). This concentration gradient may reflect intracellular or surface-bound Fog since the soluble extracellular fraction is washed away during fixation and permeabilization. To visualize the entire pool of Fog protein in living embryos (extracellular and intracellular), we engineered an endogenous Fog::SYFP2 protein fusion at the *fog* locus using CRISPR/Cas9 (referred to as Fog::YFP, Fig. S4A). Fog::YFP homozygote flies were viable and their embryos showed no defects in gastrulation (Fig. S4B,C and Movie S12). Furthermore, IHC to detect Fog::YFP with antibodies anti-GFP (to visualize the YFP tag) and anti-Fog (to visualize the Fog portion of the fusion protein) showed 1) a localization consistent with that of the known endogenous protein in both the endoderm primordium (subapical vesicular accumulation, Suppl. Fig. 4d) and the PZ (shallow apical gradient, Fig. 4D,E) and 2) a high degree of co-localization between the two detection methods. Altogether these results indicate that Fog::YFP is functional and correctly localized. In particular, they also suggest that Fog::YFP is not cleaved in the embryo despite the presence of three putative protein cleavage sites in the Fog protein sequence (*4*). To further test this latter possibility, we measured the diffusion coefficient (D) of Fog::YFP in the extracellular space using fluorescence correlation spectroscopy (FCS). Within the extracellular space between the VM and the invaginated posterior endoderm (Fig. 4F), Fog::YFP exhibited significantly slower diffusion dynamics than secreted GFP (Fig. 4G and Fig. S5A) at comparable excitation power. To circumvent artifacts induced by photophysical effects, we interpolated the values of D obtained from FCS measurements at different excitation powers (Fig. 4H and Fig. S5B). We thereby obtained for Fog::YFP a D of 55 μm^2^s^−1^, as predicted for a globular 127kDa protein diffusing in an aqueous fluid and for a secreted version of GFP a significantly higher D=87 μm^2^s^−1^ (sec::GFP, 27kDa, Fig. 4G,H and Fig. S5A, see methods). Thus, Fog::YFP is a good Fog reporter *in vivo* and we used it to image Fog distribution during wave propagation. In contrast to our expectations and the fact that intracellular Fog forms a gradient, we observed that Fog::YFP was present in the extracellular perivitelline space at uniform concentration (Movie S13). The distribution of Fog::YFP across the region of wave propagation was similar to that of sec::GFP, which reflects the available space between the apical surface of cells and the VM (Fig. 4I-K and Movies S14 and S15). Thus, endogenous Fog::YFP was uniformly distributed in the extracellular perivitelline space during wave propagation. This was independent from the presence of a maternal protein pool since a similar distribution was observed also when Fog::YFP was expressed only zygotically with a *hkb* promoter (Fig. S4E-G and Movie S16).

**Figure 4.**
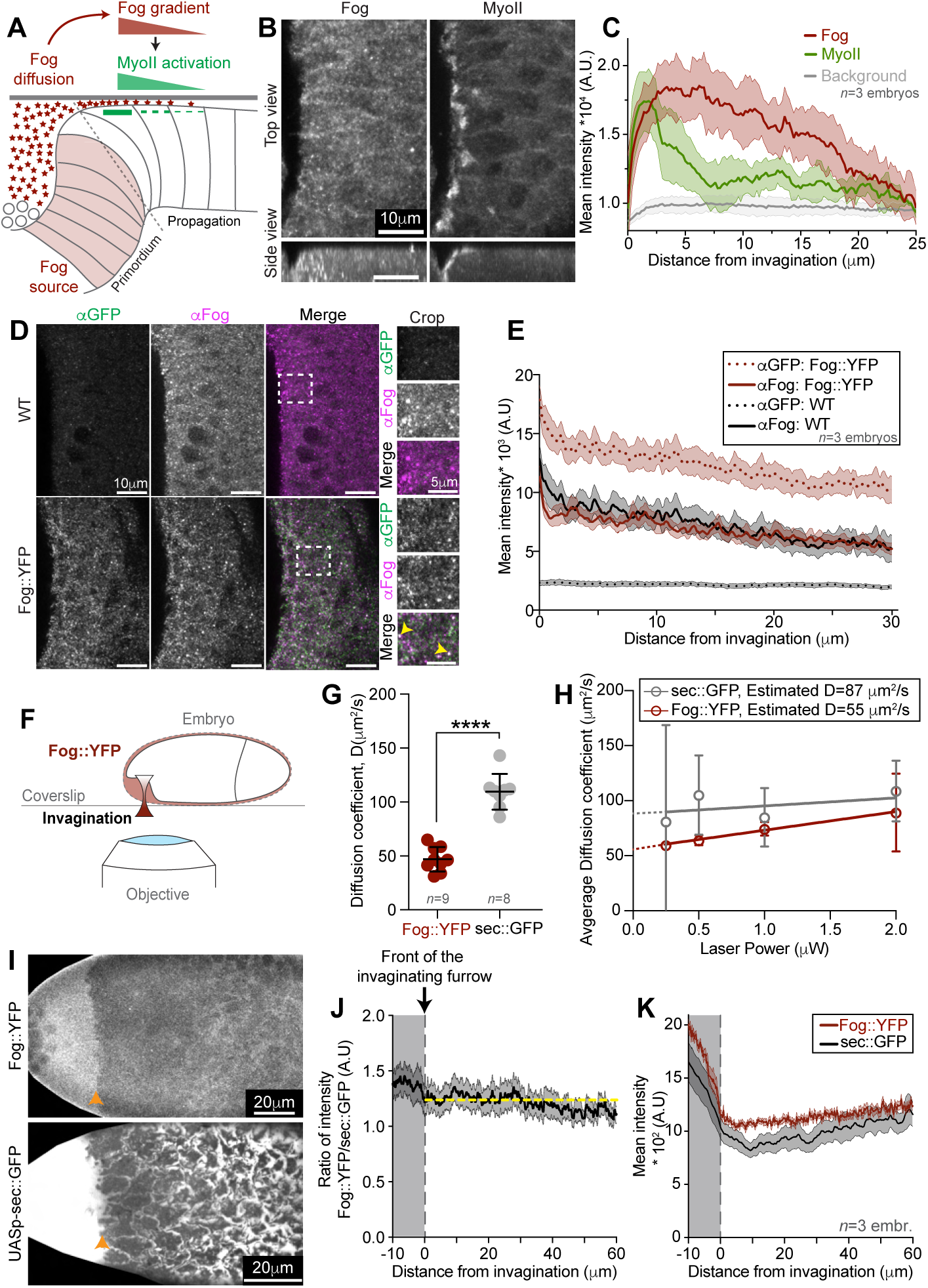
Fog::YFP does not form a concentration gradient in the extracellular space. (**A**): Illustration of the hypothesis that the gradient of MyoII (green) in the PZ depends on a concentration gradient Fog (red stars) diffusing from the invaginated primordium. (**B**): Representative confocal top and side views of Fog and MyoII during wave propagation revealed by IHC in fixed embryos. (**C**): Average spatial intensity profiles of Fog and MyoII from IHC in fixed embryos as shown in **B**. The background profile (grey) is the measurement of the embryo autofluorescence captured using 405nm laser line (see methods). *n*=3 embryos. (**D**): Representative confocal images of Fog revealed with an anti-GFP and an anti-Fog antibodies in apical sections in the dorsal-posterior region of WT (top) and Fog::YFP (bottom) embryos. (**E**): Average spatial profiles of GFP and Fog intensities from IHC in the indicated conditions. *N* =3 embryos. (**F**): Cartoon of an embryo prepared for Fluorescence correlation spectroscopy (FCS) measurement. Invagination: the FCS spot was placed in the extracellular space within the invaginating furrow. (**G**): Average diffusion coefficients of Fog::YFP and sec::GFP obtained by fitting a one-component diffusion model to FCS autocorrelation functions (ACFs) of individual measurements (*n* =9 for Fog::YFP and 8 for sec::GFP from 3 embryos each), acquired at comparable effective excitation of YFP and GFP fluorophores at 514 nm (i.e., 0.25 µW for Fog::YFP vs. 2 µW for sec::GFP). Corresponding average ACFs are shown in Suppl. Fig. 5A. \*\*\*\**P*<0.0001, unpaired *t*-test with Welch’s correction. (**H**): Quantification of the diffusion coefficients of Fog::YFP and sec::GFP from ACFs acquired at different laser powers. To cope with increased noise at low laser powers, 6-10 ACFs acquired in 3-4 embryos each were averaged per laser power and the average ACF fitted. Error bars represent 95% confidence intervals. The solid lines show a linear regression of D with laser power. The interpolation to zero excitation power (dashed lines) provides estimates for diffusion coefficients not biased by photophysical artifacts (given in the box). (**I**): Stills from time-lapse of the distribution of Fog::YFP (top) and sec::GFP (bottom) just below the vitelline membrane during wave propagation. The moving front of the invagination is marked by an orange arrowhead. (**J**): Spatial profile of the ratio of mean intensities between Fog::YFP and sec::GFP. The yellow dashed line indicates a constant ratio. The position 0μm indicates the moving front of the invagination for all time points. *n*=9 ratios from embryos. (**K**): The spatial mean intensity profiles of indicated conditions. In **J** and **K** *n*=3 embryos. Statistics: Data in **G** is mean±s.d and data in **C**, **E** and **I** are mean±s.e.m. Scale bars, 20 μm (**I**), 10 μm (**B** and **D**) and 5 μm (**D**, crop).

We conclude that extracellular Fog does not exhibit a concentration gradient in the PZ. Yet, intracellular Fog and its signalling (MyoII activation) are restricted to a dynamic domain of few cells anterior to the advancing furrow. This argues that Fog dependent signalling is organised at the cell surface in a dynamic spatial gradient, as the tissue invagination wave propagates anteriorly.

### GPCR endocytosis tunes a gradient of cell-surface bound Fog and receptor oligomerization

This led us to investigate how a MyoII activation gradient emerges in the PZ in spite of uniform extracellular Fog in the vitelline fluid. To test this, we analyzed the diffusion dynamics of Fog at the apical surface of cells in the PZ using FCS (Fig. 5A). In contrast to our measurements in the extracellular space within the posterior invagination, in the PZ, we observed two populations of Fog, characterized by two distinct diffusion coefficients (Fig. 5B,C and Fig. S5C-E). A large fraction of Fog (84% of the total pool) displayed an average diffusion coefficient D of 51±2 s.e.m μm^2^s^−1^ (D_fast_), similar to that measured in the posterior invagination and indicative of free diffusion (Fig. 5C). A minor fraction of Fog (16% of the total pool) showed much slower diffusion dynamics, with an average D of 1.6±0.2 s.e.m. μm^2^s^−1^ (D_slow_). This presumably corresponds to ligands bound to or interacting with the apical plasma membrane of cells, since it was undetectable in measurements within the extracellular space performed inside the invagination (Fig. 5B). Interestingly, the average diffusion coefficient of the Fog receptor Smog::GFP (0.54±0.04 s.e.m. μm^2^s^− 1^) at the apical cell surface in the PZ measured by FCS was in the same range, but nonetheless significantly lower than D_slow_ of Fog (Fig. 5D). This suggests that the slow-diffusing fraction of Fog (Fog_slow_) could correspond to ligands transiently bound to their receptor in the PZ. To further test this, we sought to increase the half-life of the receptor at the surface by inhibiting GCPR endocytosis (*gprk2* RNAi) and measured Fog_slow_ in the PZ. We found that in *gprk2* RNAi embryos the Fog_slow_ fraction increased to 30±15% (mean±s.e.m.) of the total pool from 16±1% (mean±s.e.m.) in WT, consistent with the idea that Fog_slow_ is receptor-bound on the surface of cells in the PZ (Fig. 5B,D). We next examined whether Fog_slow_ forms a spatial gradient in the PZ and we found that in WT embryos Fog_slow_ is higher close to the invagination, and decreases gradually at increasing distance from the invaginating furrow (Fig. 5E,F, gradient slope: −1.4*10^−3^ μm^−1^). Furthermore, blocking GPCR endocytosis not only globally increased Fog_slow_ but also reduced the slope of its gradient (Fig. 5G,H, black bold line, gradient slope: −2.6*10^−4^ μm^−1^). These results are consistent with the observed higher levels of MyoII and its expanded gradient in *gprk2* RNAi embryos and suggest that the Fog activity gradient might emerge from a gradient of ligand-receptor binding within the PZ. Next, we sought to measure Fog receptor activation within the PZ to see if that also formed a gradient as predicted from the distribution of Fog_slow_. GPCR ligand binding and activation is known to induce receptor clustering (oligomerization) (*37, 40, 41*), which can be measured by FCS (*37*). Using FCS, we measured the per-particle brightness, which reflects the degree of molecular oligomerization (see methods), of Smog::GFP on the cell apical surface in the PZ. Consistent with previous studies (*37*), we found that the molecular brightness of Smog::GFP decreased upon depletion of the ligand Fog (Fig. 5i) and increased upon depletion of Gprk2 (Fig. 5J) without changing its diffusion coefficient and only slightly affecting its total concentration (Fig. S6A-C). We next compared molecular brightness of Smog::GFP with that of the known trimeric protein VSV-G::GFP and found that the brightness of VSV-G::GFP was significantly higher than that of Smog::GFP in both the PZ (Fig. 5I,J) and the lateral ectoderm (Fig. 6D). Furthermore, since VSV-G::GFP brightness was not affected by neither Fog nor Gprk2 depletion (Fig. 5I,J), these results confirmed that the observed effects on Smog::GFP are receptor specific and that its oligomerization is a proxy for GPCR activation.

**Figure 5.**
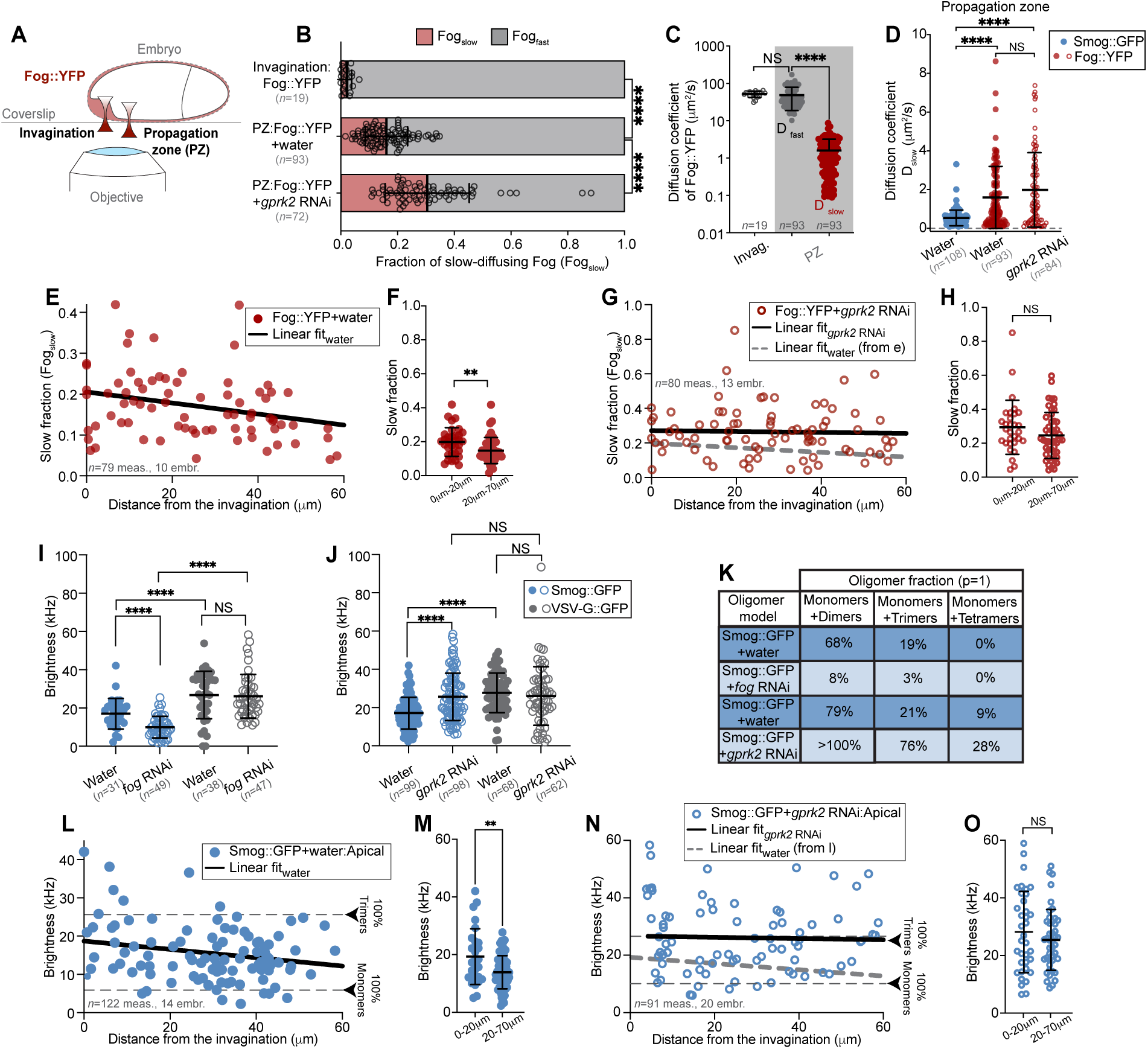
Slowly diffusing Fog::YFP and Smog::GFP homoclusters form a gradient at the apical cell surface in the PZ. (**A**): Cartoon of an embryo prepared for fluorescence correlation spectroscopy (FCS) measurement. Invagination: FCS spot was placed in the extracellular space within the invaginating furrow. Propagation zone (PZ): FCS spot was placed on the apical surface or at junctions during wave propagation. (**B**): Fraction of slow-(Fog_slow_) and fast-diffusing (Fog_fast_) pools of Fog::YFP in indicated conditions, obtained from two-component FCS analysis. Fog_fast_+Fog_slow_=1. *n=*19 measurements from 11 embryos (invagination Fog::YFP), 93 measurements from 11 embryos (PZ Fog::YFP + water) and 72 measurements from 12 embryos (PZ Fog::YFP + *gprk2* RNAi). (**C**): Measurements of the diffusion coefficient (D) of Fog::YFP in the invagination (white area) or in the PZ (grey area). In the invagination D was measured by fitting ACFs with a one-component diffusion model (as in Fig. S5C), in the PZ the diffusion coefficients of the slow-(D_slow_) and fast-diffusing (D_fast_) fractions of Fog::YFP were obtained by fitting ACFs with a two-component diffusion model (as in Fig. S5E). *n=*19 measurements for the invagination, and 93 measurements for both D_slow_ and D_fast_ from 8 embryos each. NS, *P*= 0.4253, \*\*\*\**P*<0.0001, unpaired *t*-test with Welch’s correction. (**D**): Measured diffusion coefficients of the slow fraction in the PZ for the indicated molecules in the indicated conditions. *n=* 108 measurements from 6 embryos (Smog::GFP + water), 93 measurements from 8 embryos (Fog::YFP + Water) and 84 measurements from 12 embryos (Fog::YFP + *gprk2* RNAi). NS, *P*= 0.1464, \*\*\*\**P*<0.0001, unpaired *t*-test with Welch’s correction. (**E-H**): Measurements of the fraction of slow-diffusing Fog (Fog_slow_) at different distances from the invaginating furrow during wave propagation in water (**E**, *n=* 79 measurements from 10 embryos) and *gprk2* RNAi (**G**, *n=*80 measurements from 13 embryos) injected embryos. The solid black lines represent simple linear regression fits. The dashed grey line in **G** represents the simple linear regression fit from **e** for comparison. **F** and **H,** Fog_slow_ for the indicated bins of distance in water (**F**, *n=*79 measurements from 10 embryos, \*\**P*=0.0072) and *gprk2* RNAi (**H**, *n=*80 measurements from 13 embryos, NS, *P*=0.1722) injected embryos. (**I-J**): Measurements of Smog::GFP (blue) and VSV-G::GFP (grey) molecular brightness (degree of oligomerization) measured at the apical surface of cells in the PZ during wave propagation in the indicated conditions. In **I**, *n=*31 measurements from 8 embryos (Smog::GFP + water), 49 measurements from 10 embryos (Smog::GFP + *fog* RNAi), 38 measurements from 5 embryos (VSV-G::GFP + water) and 47 measurements from 6 embryos (VSV-G::GFP + *fog*RNAi). NS, *P*= 0.3970, \*\*\*\**P*<0.0001, unpaired *t*-test with Welch’s correction. In **J**, *n=*99 measurements from 14 embryos (Smog::GFP + water), 98 measurements from 20 embryos (Smog::GFP + *gprk2*RNAi), 68 measurements from 5 embryos (VSV-G::GFP + water) and 62 measurements from 8 embryos (VSV-G::GFP + *gprk2* RNAi). NS, *P*= 0.4772 (VSV-G::GFP + water vs VSV-G::GFP + *gprk2* RNAi), NS, *P*= 0.8437 (Smog::GFP + *gprk2* RNAi vs VSV-G::GFP + *gprk2* RNAi), \*\*\*\**P*<0.0001, unpaired *t*-test with Welch’s correction. (**K**): Estimation of the fraction of monomers and higher-order oligomers (see methods) of Smog::GFP at the apical surface of cells in the PZ during wave propagation in indicated conditions. ‘p’ is the probability of each GFP label being fluorescent (see methods and Fig. S6E for p=0.7). (**L**) and (**N**): Measurements of apical Smog::GFP molecular brightness at different distances from the invaginating furrow during wave propagation in water (**L**, *n=*122 measurements from 14 embryos) and *gprk2* RNAi (**N**, *n=* 91 measurements from 20 embryos) injected embryos. The dashed gray lines represent the estimated brightness of a scenario with 100% monomers (bottom line) and 100% trimers (top line). Solid black lines represent simple linear regression fitting. The dashed thick grey line in **N** represents the simple linear regression fit from **L** for comparison. (**M**) and (**O**): Apical Smog::GFP molecular brightness for the indicated bins of distance in water (**M**, *n=*122 measurements from 14 embryos, \*\**P*=0.0088) and *gprk2* RNAi (**O**, *n=*91 measurements from 20 embryos, NS,*P*=0.3373) injected embryos. Statistics: Data are mean±s.d.

**Figure 6.**
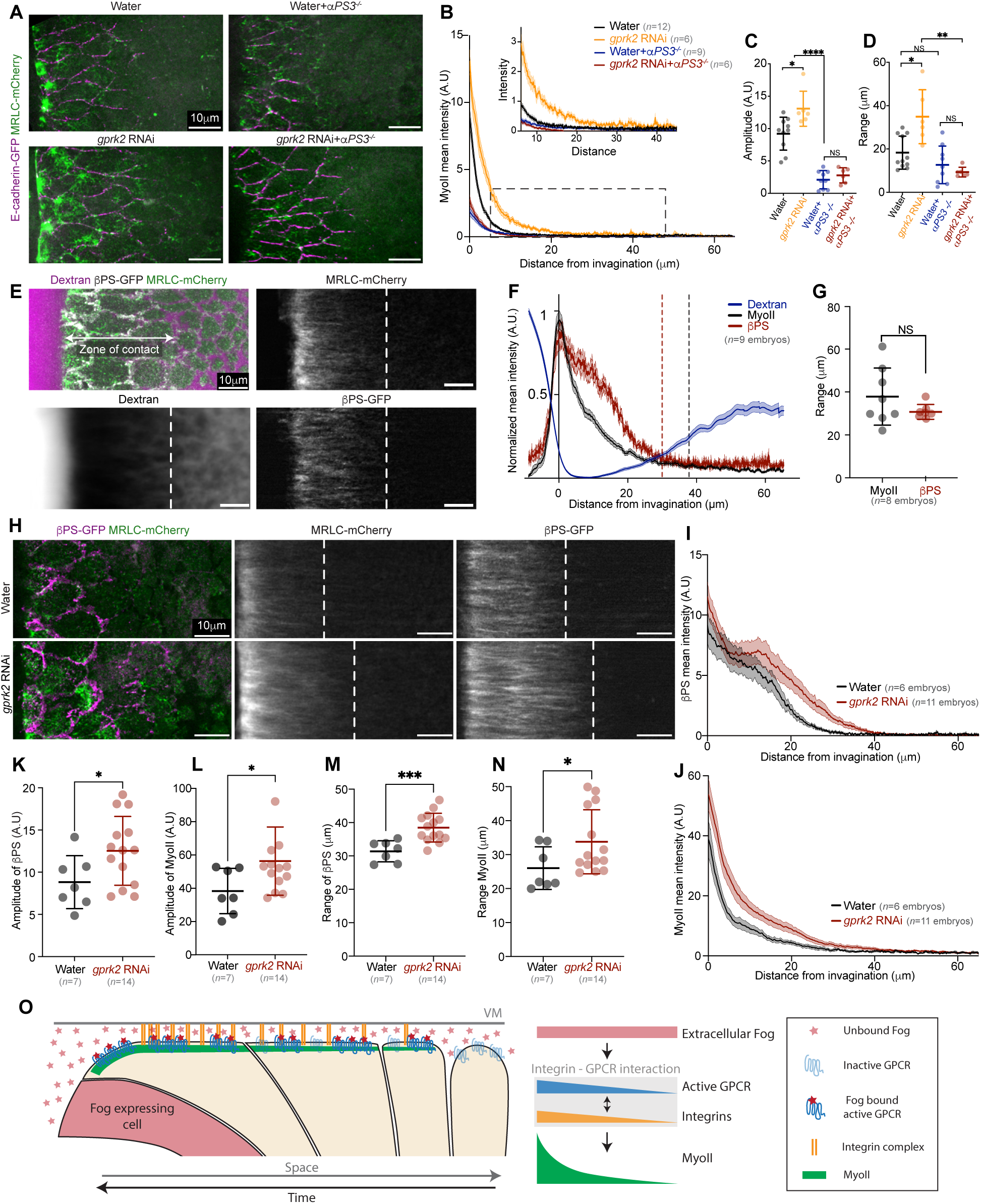
Integrin activation upon contact with the VM define range and activity of the Fog signalling gradient. (**A**): Stills from time-lapses of MyoII in the PZ in control (Water), integrins mutant (Water + *αPS3^−/−^*), and *gprk2* RNAi injections in WT (*gprk2* RNAi) or integrin mutant embryos (*gprk2* RNAi *+ αPS3^−/−^*). (**B**): Average spatial profiles of MyoII mean intensity in the indicated conditions. The inset is a zoomed view of the dashed boxed region. (**C-D**): Measurements of the amplitude (**C**) and range (**D**) of the MyoII gradients in the indicated conditions. *n*=12 embryos (Water), 8 embryos (*gprk2* RNAi), 9 embryos (water + *αPS3^−/−^*) and 6 embryos (*gprk2* RNAi + *αPS3^−/−^*). In **C,** \**P*=0.0177, \*\*\*\**P*<0.0001 and NS *P*=0.3239, unpaired *t*-test with Welch’s correction. In **D,** \**P*=0.0204, \*\**P*=0.0029, NS *P*=0.2973 (Water+*αPS3^−/−^ vs Gprk2*RNAi*+αPS3^−/−^)* and NS, *P*=0.1476 (Water *vs αPS3^−/−^)* unpaired *t*-test with Welch’s correction. (**E**): On the top left, a still from a timelapse of MyoII (green) and βPS integrin (white) in embryos injected with dextran to label the extracellular space (magenta). The other images are temporal average projections in the referential of the moving invagination (see methods) of MyoII (top right), dextran (bottom left) and βPS::GFP (bottom right). The white dashed line represents the zone where the tissue is in contact with the VM. (**F**): Average normalized spatial profiles of the mean intensity of MyoII (black), βPS::GFP (red) and dextran (blue). The black and red dashed lines indicate the range of MyoII activation and βPS::GFP focal complex formation respectively, estimated as indicated in the methods. *n*=9 embryos. (**G**): Measurements of the range of MyoII activation and βPS::GFP focal complex formation in individual embryos. *n*=8 embryos each. *P*=0.0911, unpaired *t*-test with Welch’s correction. (**H**): Left: High-resolution images of MyoII and βPS::GFP in the PZ in control (water, top) and *gprk2* RNAi injected (bottom) embryos during wave propagation. Centre and right: Temporal average projection of MyoII and βPS::GFP in the indicated conditions. The dashed white line represents the zone where the tissue is in contact with the VM. *n*=6 embryos (Water) and 11 embryos (*gprk2* RNAi). (**I-J**): Average mean intensity profiles of βPS (**I**) and MyoII (**J**) in the PZ in the indicated conditions. *n*=6 embryos (Water) and 11 embryos (*gprk2* RNAi). (**K-N**): Measurements of the intensity profile amplitude (**K**-**L**) and the range of MyoII activation/βPS focal complex formation (**M**-**N**) in indicated conditions. *n*=7 embryos (Water) and 14 embryos (*gprk2* RNAi). In **K,** \**P*=0.0353, **L,** \**P*=0.0285, **M,** \*\*\**P*=0.0006, **N,** \**P*=0.0381 using an unpaired *t*-test with Welch’s correction. **O**, Cartoon model of the gradient of MyoII activation during wave propagation from extracellular Fog acting as a mechanogen. One Fog expressing cell within the primordium (Fog source) is labelled in red. Unbound extracellular Fog and inactive GPCR receptors are semi-transparent. The active GPCR bound to Fog are in solid colors. The zone of contact of the tissue with the vitelline membrane is labelled by the presence of bound integrin complexes. Statistics: Data in **C**, **D**, **G**, **K**, **L**, **M**, **N** are mean±s.d. Data in **B**, **F**, **I** and **J** are mean±s.e.m. Scale bars, 10 μm.

The comparison of molecular brightness with VSV-G::GFP also suggests that, in the simplest scenario, Smog::GFP is present as a mixture of dimers and monomers in WT embryos (molecular brightness lower than VSV-G::GFP), while oligomerization increases upon blocking receptor endocytosis (similar brightness of the trimeric protein VSV-G::GFP, Fig. 5J) and decreases in the absence of the ligand Fog during wave propagation (Fig. 5I). Since our data indicate that Smog::GFP exists as monomers and higher-order oligomers (dimers, trimers, etc.), we next estimated the fraction of oligomers from the measured brightness, assuming a linear dependence of the brightness on the number of protomers (see methods). Since only the average brightness can be directly extracted from FCS measurements, the exact type of oligomeric species cannot be determined. Nevertheless, for a particular model, i.e. a two-species mixture of monomers and oligomers, the fraction of monomers and oligomers can be estimated (see methods). We calculated these numbers for several possible scenarios (monomers with either dimers, trimers or tetramers) in WT, *fog* and *gprk2* RNAi embryos taking previously reported photophysical properties of GFP (*42*) into account (Fig. 5K and Fig. S6E). Depleting the ligand Fog by RNAi reduced the probability of higher-order oligomer formation for Smog::GFP by ∼8 folds, i.e. from ∼68% dimers in WT embryos to less than 8% in *fog* RNAi embryos (Fig. 5K and Fig. S6E). Conversely, blocking receptor endocytosis significantly increased oligomer formation by ∼21% when considering the couple dimers/monomers (from 79% to 100%, Fig. 5K and Fig. S6E) or by ∼55% when considering the couple trimers/monomers (from 21% to 76%, Fig. 5K and Fig. 6E). To test whether Smog::GFP oligomers are distributed in a gradient in the PZ, we measured Smog::GFP brightness on the apical cell surface and at lateral junctions as a function of the distance from the invaginating furrow in the PZ (Fig. 5L,M and Fig. 6F,G). In WT embryos we found higher Smog::GFP brightness closer to the invagination (0-20μm) than at larger distances (20-70μm), with similar trends both on the apical surface and at junctions (Fig. 5L,M, gradient slope: −0.11 μm^−1^ in the apical region and Fig. S6F,G, gradient slope −4.5*10^−2^ μm^−1^ in the junctional region). *gprk2* RNAi reduced this spatial difference by increasing Smog::GFP brightness both close and at a distance from the invagination (Fig. 5N,O, gradient slope: −2.2*10^−2^ μm^−1^ in the apical region and Fig. S6H,I, gradient slope: 2.9*10^−4^ μm^− 1^ in the junctional region), indicating that the gradient of Smog::GFP oligomerization depends on GPCR endocytosis. Importantly, this spatial dependence of oligomerization was specific to Smog::GFP as no gradient (apical region) or even an inverse gradient (at junctions) was observed in VSV-G::GFP control embryos (Fig. S6J-M, gradient slope: −0.5*10^−1^ μm^−1^ in the apical region and 0.7*10^−1^ μm^−1^ in the junctional region).

Altogether, our data indicate that Fog binding to the cell apical surface induces a gradient of receptor activation and oligomerization, which is tuned by receptor endocytosis.

### Integrin activation by contact with the vitelline membrane tunes the Fog activity gradient

The gradient of GPCR activation and oligomerization tunes the multicellular spatial pattern of MyoII activation during wave propagation. Integrins, which are activated when cells of the PZ contact the VM, are also involved in MyoII activation (*6, 30*). We showed earlier that the *⍺*PS3 integrin Scab amplifies MyoII levels in a positive feedback mechanism (*30*). Consistent with this, in *αPS3* null mutants (*αPS3^−/−^*), MyoII was activated at much lower levels than in controls, although higher than those observed in the absence of G*⍺* signalling (Fig. S7A,B). Moreover, in these mutants MyoII activation in the PZ still followed an exponential gradient with similar range than WT, albeit with lower amplitude, which led to an increased length scale compared to WT embryos (Fig. 6A-D and Fig. S7C-E and Movie S17). This suggests that amplification of MyoII activation by integrins is not uniform in the PZ (uniform amplification by a constant factor would not change the gradient length scale). This led us to address whether integrins only amplify MyoII activation in the cell near the edge of invagination front, as previously suggested (*30*), or whether they also have a role in regulating the spatial range of the multicellular Fog/GPCR activity gradient. Since amplitude, range and length scale of the gradient are increased significantly when GPCR endocytosis is inhibited in *gprk2* RNAi embryos (Fig. 3G-I, and Movie S11), we tested whether integrins are required for this effect and thus for the process of gradient formation by acting everywhere in the field where Fog and GPCR signalling can be active. Strikingly, removing integrins in embryos depleted for Gprk2 (*gprk2*RNAi + *αPS3^−/−^*) dramatically reduced both the levels and the range of MyoII to levels similar to those of the knock-out of integrins (*αPS3^−/−^*, Fig. 6A-D, Fig. S7c and Movie S17). These results show that integrins are required for both the expanded gradient and the higher levels of MyoII observed following the hyperactivation of Fog-GPCR signalling. Additionally, it shows that blocking endocytosis has no impact on the shape of the gradient if integrins are absent. We conclude that integrins are part of the multicellular gradient formation mechanism and do not simply amplify MyoII activation in the cell near the invagination front.

Integrins accumulate and form bright puncta in the PZ upon apical cell contact with the overlying vitelline membrane. The range of this contact might thus contribute to determining the range of the gradient of MyoII and GPCR signalling. Consistent with this idea, integrin puncta, visualized with the endogenously GFP-tagged βPS integrin Myospheroid (βPS::GFP hereafter), accumulated at highest levels close to the invaginating furrow and decreased more anteriorly forming a gradient with a range identical to that of MyoII activation in the PZ (Fig. 6E-G, and Fig. S7F). This integrin gradient formed within the region of contact of the tissue with the VM, as revealed by the exclusion of dextran injected in the perivitelline space from the region where βPS::GFP forms bright puncta (Fig. 6E-G). Furthermore, the depletion of Gprk2 increased the amplitude and the range of the gradient of βPS::GFP (Fig. 6H-N and Movie S18) as well as the range of tissue contact with the VM (monitored by the exclusion of dextran and by the distance of E-cad junction from the VM, Fig. S7G-K and Movie S19).

Together we conclude that during wave propagation MyoII is activated where integrins are in contact with the VM and that the range of the exponential gradient of MyoII activation during wave propagation depends on the contact of the tissue with the VM in a manner that involves integrin activation and their cross-talk with Fog-GPCR signalling.

## Discussion

Cell and tissue mechanics must be regulated in space and time to drive tissue morphogenesis. The mechanisms of such regulation remain poorly understood. Here, we report how a dynamic gradient of cell contractility essential for *Drosophila* embryo gastrulation is set up by direct, long-range and concentration-dependent signalling of a diffusing ligand (Fog) of a GPCR (Smog). This is consistent with a hypothesis referred to as a mechanogen (*2, 3*) where, by analogy to morphogens that spatially organise cell identities transcriptionally, diffusible molecules directly pattern, through activity gradients, mechanical properties (e.g. contractility) in fields of cells. Notably, this hypothesis contrasts with classical mechanisms where mechanical regulation is organized in two tiers and effected downstream of spatial patterning of cell identities via transcriptional regulation. For instance, the morphogen Dorsal activates *twist* and *snail,* which define the mesodermal identity of ventral cells, in *Drosophila* embryos and in turn regulate transcriptionally the GPCR ligand Fog to induce their tissue autonomous apical constriction and tissue invagination. Here we show that Fog also acts tissue non-autonomously to control a gradient of MyoII activation required for a wave of tissue invagination driving the morphogenesis of the posterior endoderm. The shape of this gradient is important as shown by the fact that in mutant conditions where the amplitude and the range of MyoII activation increase (e.g. *hkb-fog* or *gprk2* RNAi, Fig. 3) the speed of the tissue propagation wave was reduced, due to increased adhesion to the vitelline membrane and reduced de-adhesion rate (*30*).

We demonstrate that Fog produced zygotically in the posterior endoderm primordium, not only drives apical constriction and tissue invagination cell autonomously, but it also diffuses in the extracellular perivitelline space to control an exponential gradient of MyoII activation more anteriorly in cells of the propagation zone. Thus, while Fog has long been proposed to act as a switch factor to locally convert a genetic pattern into a pattern of cortical mechanics driving tissue dynamics, we demonstrate that it is also able to diffuse and pattern cortical mechanics at a distance in the zone of wave propagation. Similar to morphogens, we show that the amplitude and the range of Fog activity (MyoII activation) depend on the rate of ligand production at the source (the primordium) and the rate of its clearance by endocytosis in the target tissue. Thus, since Fog diffuses from a localized source and directly tunes tissue mechanical properties (actomyosin contractility) in a concentration-dependent manner, it fulfils the proposed definition of a mechanogen (*2, 3*). Strikingly, however, the activity pattern of Fog does not reflect its protein distribution in the extracellular space where it is distributed uniformly. Yet, we found that Fog forms a gradient of cell surface-bound ligand (Fig. 5E) associated with a gradient of receptor oligomerization (Fig. 5L). Furthermore, both the gradient of surface-bound Fog and receptor oligomers are regulated by GPCR endocytosis (Fig. 5G,N). When GPCR endocytosis is blocked GPCR oligomerization no longer forms a gradient (Fig. 5N) and the range of the contractility gradient is expanded towards the anterior (Fig. 3G,H). It is possible that the gradient might also require extracellular regulation of Fog protein activity by proteins capable of binding and/or activating/inactivating Fog or its receptor at the cell surface. The formation of a Dpp morphogen gradient in the vitelline space requires binding to its inhibitor Sog and a protease Tld that enables Dpp activity in a narrower dorsal domain than where Dpp is expressed and diffuses (*43, 44*). Post-translational modification by an extracellular inhibitor may also spatially restrict Fog activity and thereby be part of the gradient formation mechanism.

Another salient feature of the Fog activity gradient on the cell surface is that it translocates with the front of tissue invagination as it sweeps as a wave towards the anterior of the embryo. As a result, the gradient is constantly renewed over time. Because of the wave nature of the process, any cell within the field of the gradient moves towards the invagination front and experiences a temporal increase in MyoII activation (Fig 1J, and Fig. S7B). In other words, the spatial nature of the gradient in the referential of the embryo is inherently associated with a temporal “maturation” of cell signalling in the referential of the cells. Therefore, the mechanogen gradient reported here is coupled to a temporal integration process that involves GPCR activation by Fog and integrins (Fig. 6O).

The Fog activity gradient self-organises at the cell surface while Fog is at a uniform concentration in the vitelline fluid. Since tissue deformation during the wave of invagination is expected to cause turbulent hydrodynamic flow in the vitelline fluid, which could disturb extracellular fluid gradients, the ability to self-organize a Fog surface activity gradient at the cell surface via receptor endocytosis and integrins (see below) may be an adaptive mechanism to ensure robust gradient formation.

Finally, we report here that integrins control Fog-GPCR signalling during wave propagation and thereby shape its activity gradient. In the absence of integrins blocking GPCR endocytosis (*gprk2* RNAi) no longer extends the Fog activity gradient and MyoII activation, indicating that integrins are required for robust Fog-GPCR signalling. Integrin-based adhesion may inhibit endocytosis of the GPCR Smog (e.g. by frustrated endocytosis (*45*)), promote GPCR oligomerization, potentiate binding of Fog to the GPCR or other downstream molecular effector. The reported function of integrins in setting up the Fog activity gradient is interesting because during wave propagation integrins are activated by the initial posterior tissue invagination which brings more anterior cells in contact with the vitelline membrane. Within this region of contact integrins form a gradient due to a positive feedback regulation with MyoII activation (*30*). Thus, the amplitude and the length scale of the biochemical gradient that activates MyoII during wave propagation require feedback regulation by integrins which itself rests on the geometry of tissue invagination and interaction with the vitelline membrane. Such feedback loops tune the length scale of the mechanogen gradient reported here. This work exemplifies the rich modalities of spatial patterning of cell and tissue mechanics during development independent of gene transcriptional regulation. It will be especially intriguing to consider in the future how mechanical patterning by biochemical signalling may depend on the mechanical and geometric properties of the tissues where this is taking place.

## Author contributions

C.C. and T.L. conceived and supervised the study. G.M., C.C. and T.L. planned the experiments with help from V.D-E. for the planning of the FCS experiments. G.M. performed all the experiments and quantifications except for those in Fig. 6E-J, Fig. S3E and Fig. S7F-I, which were performed by C.C. V.D-E. analyzed and quantified all the FCS data. J.M.P. and E.D.S designed and generated all the molecular constructs with the help of G.M. All authors discussed the data. G.M. and C.C. prepared the figures. G.M., C.C. and T.L. wrote the paper with help from V.D-E. for the discussion of the FCS results. All authors made comments.

## Acknowledgements

We thank all members of the Lecuit lab for stimulating discussion and feedback during the course of this project. We thank the imaging facility at IBDM, member of the National Infrastructure France-BioImaging (https://ror.org/01y7vt929) supported by the French National Research Agency (ANR-24-INBS-0005 FBI BIOGEN), for assistance with maintenance of the microscopes; FlyBase for maintaining curated databases; and Bloomington Drosophila Stock Center for providing fly stocks.

## Funding

This work was supported by the ERC grant SelfControl #788308. C.C. is supported by the CNRS; T.L.is supported by the Collège de France; G.M. was supported by the ERC grant SelfControl #788308 and an ATER fellowship from the Collège de France. V.D-E. was supported by Human Frontier Science Program Long-term Fellowship (HFSP LT0058/2022-L).

---

**Figure S1.**
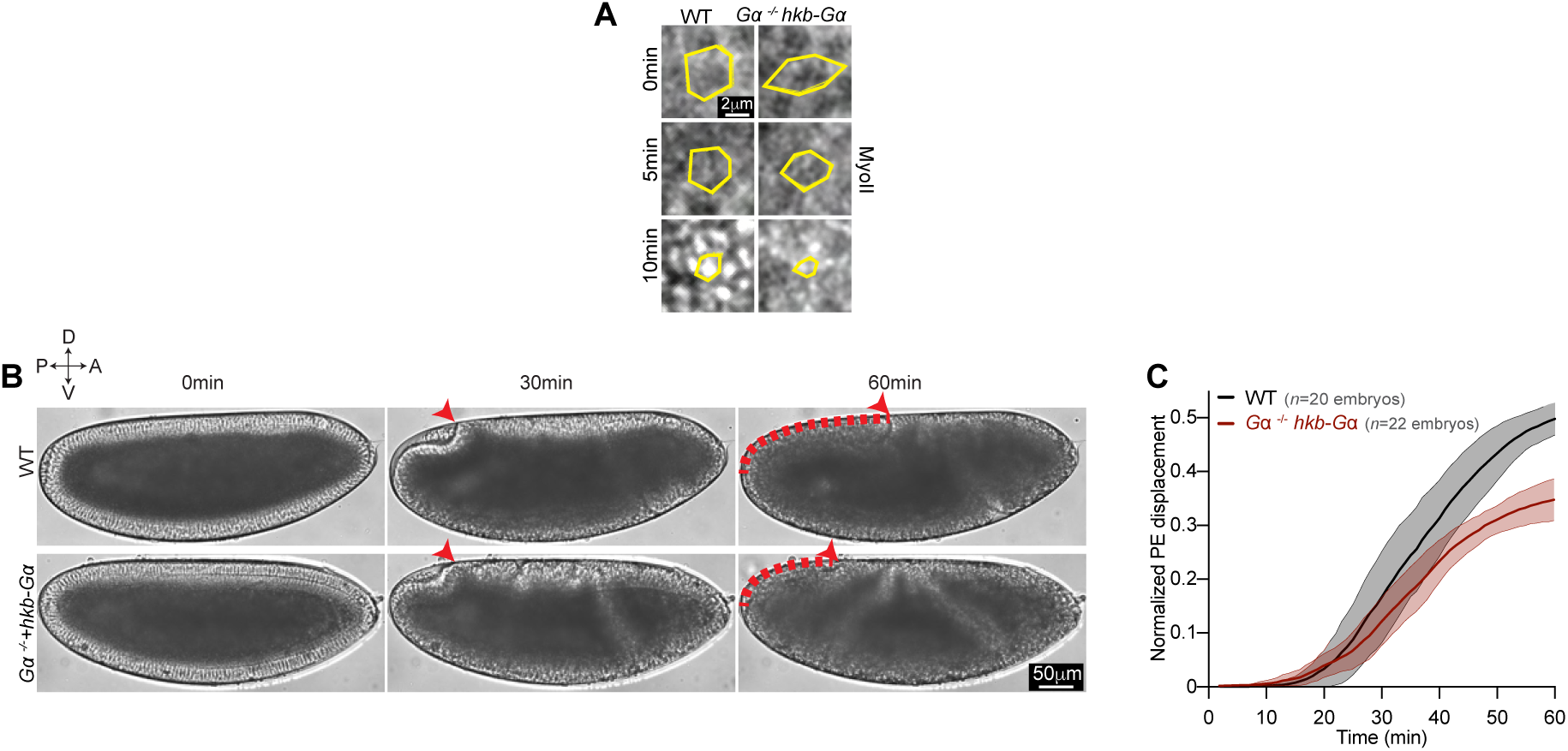
The anterior movement of the posterior endoderm is defective in Gα^−/−^*hkb*-Gα embryos. (**A**): Close-up views over time of MyoII of the cells in the primordium region depicted in 1e. In yellow the contours of a cell tracked over time. (**B**): Stills from brightfield time-lapses in the indicated conditions. In *Gα*^−/−^*hkb*-Gα embryos the primordium invaginates normally (time 30 min) but the subsequent movement towards the anterior fails (time 60min). The red arrowheads mark the dorsal-anterior edge of the invagination and red dashed lines indicate the displacement of the posterior endoderm at the indicated time. (**C**): Quantifications of the normalized posterior endoderm (PE) displacement over time in WT and *Gα*^−/−^ *hkb*-Gα embryos. *n=* 20 and 22 embryos for control and *Gα*^−/−^ *hkb*-Gα embryos respectively. Time 0 is the time when the cellularization front reaches the basal side of nuclei on the dorsal side of the embryo. Statistics: in **C** traces are mean±s.d. Scale bars, 50 μm (**B**) and 5 μm (**A**).

**Figure S2.**
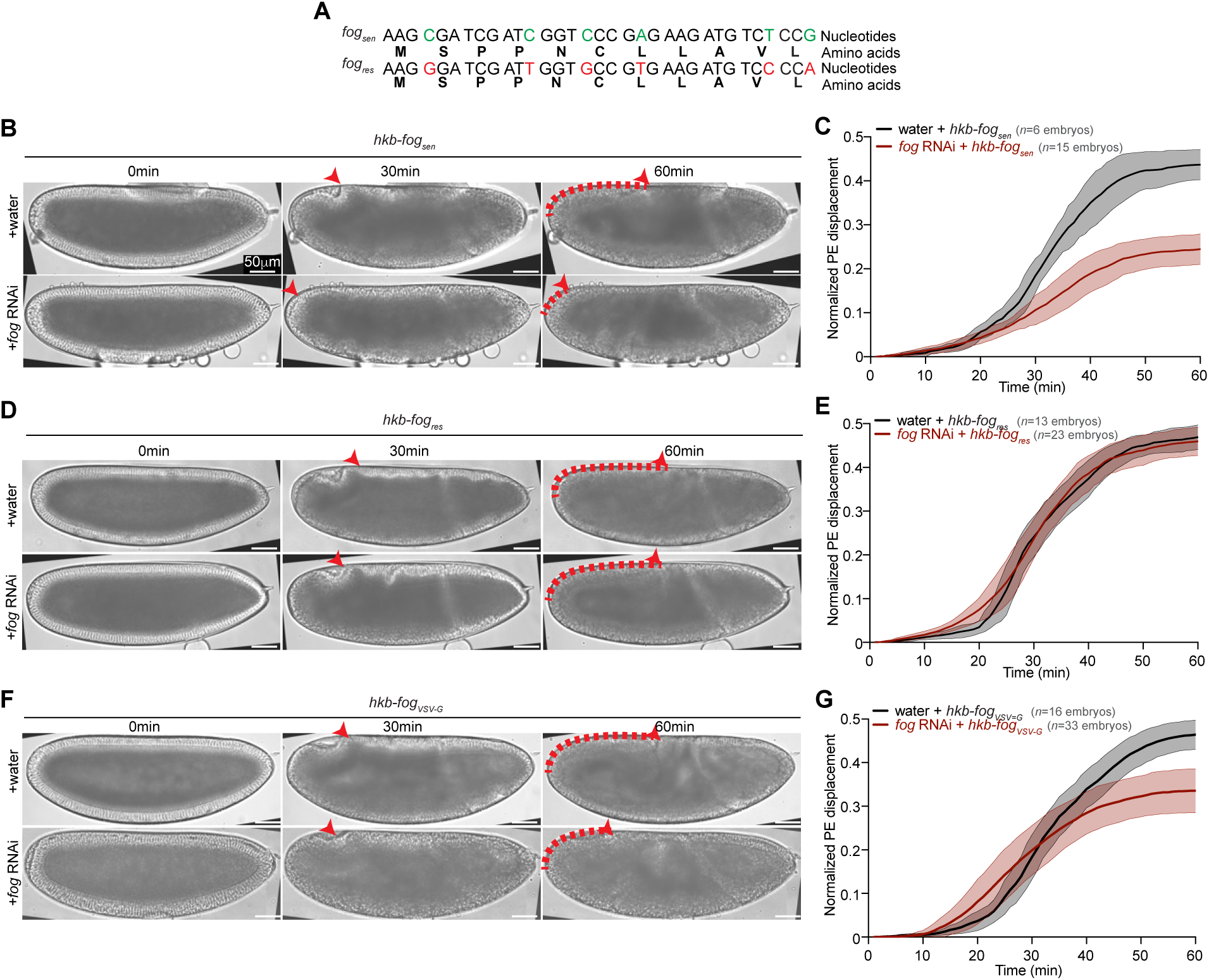
Tethering Fog to the membrane impairs the anterior movement of the posterior endoderm. (**A**) Alignment of a small portion of the nucleotide sequences of the original (RNAi-sensitive, *fog_sen_*) and the RNAi-resistant (*fog_res_*) versions of *fog*. Since the entire sequence targeted by *fog* dsRNAs was modified using codon redundancy, it is possible to obtain an RNAi-resistant version of the *fog* mRNA. (**B**): Stills from brightfield time-lapses of embryos in *hkb-fog_sen_* embryos injected with water (control) or *fog* RNAi. Red arrowheads mark the dorsal edge of the endoderm invagination and red dashed lines indicate the displacement of the posterior endoderm at the indicated times. (**C**): Quantifications of the normalized posterior endoderm (PE) displacement over time in the indicated conditions. *n=*6 (*hkb-fog_sen_* + water) and 15 (*hkb-fog_sen_* + *fog* RNAi) embryos respectively. (**D**): Stills from brightfield time-lapses of embryos in *hkb-fog_res_* embryos injected with water (control) or *fog* RNAi. (**E)**: Quantification of the normalized posterior endoderm (PE) displacement over time in the indicated conditions. *N=*13 (*hkb-fog_res_* + water) and 23 (*hkb-fog_res_* + fog RNAi) embryos. (**F**): Stills from brightfield time-lapses of embryos in *hkb-fog_VSV-G_* embryos injected with water (control) or *fog* RNAi. (**G**): Quantification of the normalized posterior endoderm (PE) displacement over time in the indicated conditions. *n=*16 (*hkb-fog_VSV-G_* + water) and 33 (*hkb-fog_VSV-G_* + *fog* RNAi) embryos. Time 0 is the time when the cellularization front reaches the basal side of nuclei on the dorsal side of the embryo. Statistics: Data in **C**, **E** and **G** are mean±s.d. Scale bars, 50 μm.

**Figure S3.**
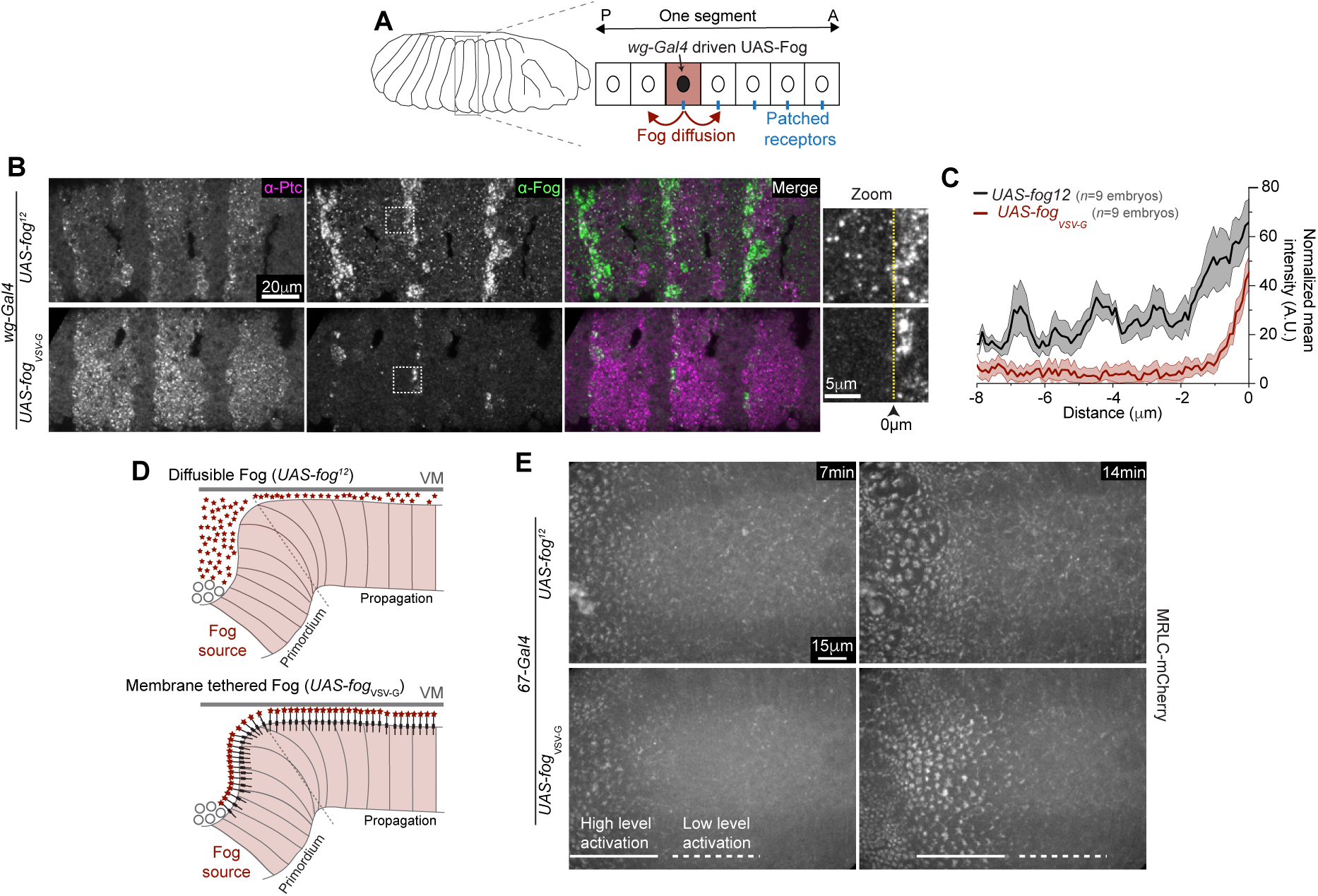
Membrane-tethered Fog is functional and activates MyoII. (**A**): Schematic of the striped expression using the *wingless* Gal4 (*wg-Gal4*) driver to express a diffusible (*UAS-fog^12^*) and a membrane-tethered (*UAS-fog_VSV-G_*) Fog at embryonic developmental stage 12. Patched (Ptc) marks the posterior edge of *wg* expression within each segment. (**B**): Representative confocal images of Ptc (magenta) and Fog (green) in embryos expressing diffusible (top, *UAS-fog^12^*) or membrane-tethered Fog (bottom, *UAS-fog_VSV-G_*). On the right, a zoomed view of Fog around the *wingless* stripe. The yellow dotted line indicates the border of the stripe and was defined as the origin for intensity profile quantifications in **C**. (**C**): Spatial profiles of Fog mean intensity in the indicated conditions. Intensity values are normalized to maximum levels for visualization purposes. *n=*1 segment from 9 embryos each. (**D**): Cartoon of side views of the posterior endoderm in embryos homogenously expressing a diffusible (*UAS-fog^12^*) or a membrane-tethered (*UAS-fog_VSV-G_*) version of Fog. (**E**): Representative stills over time of MyoII patterns in the dorsal epithelium in embryos in the indicated conditions. *n* =3 embryos each. Statistics: Data in **C** is mean±s.e.m. Scale bars, 20 μm (**B**), 15 μm (**E**) and 5 μm (**B**, zoom).

**Figure S4.**
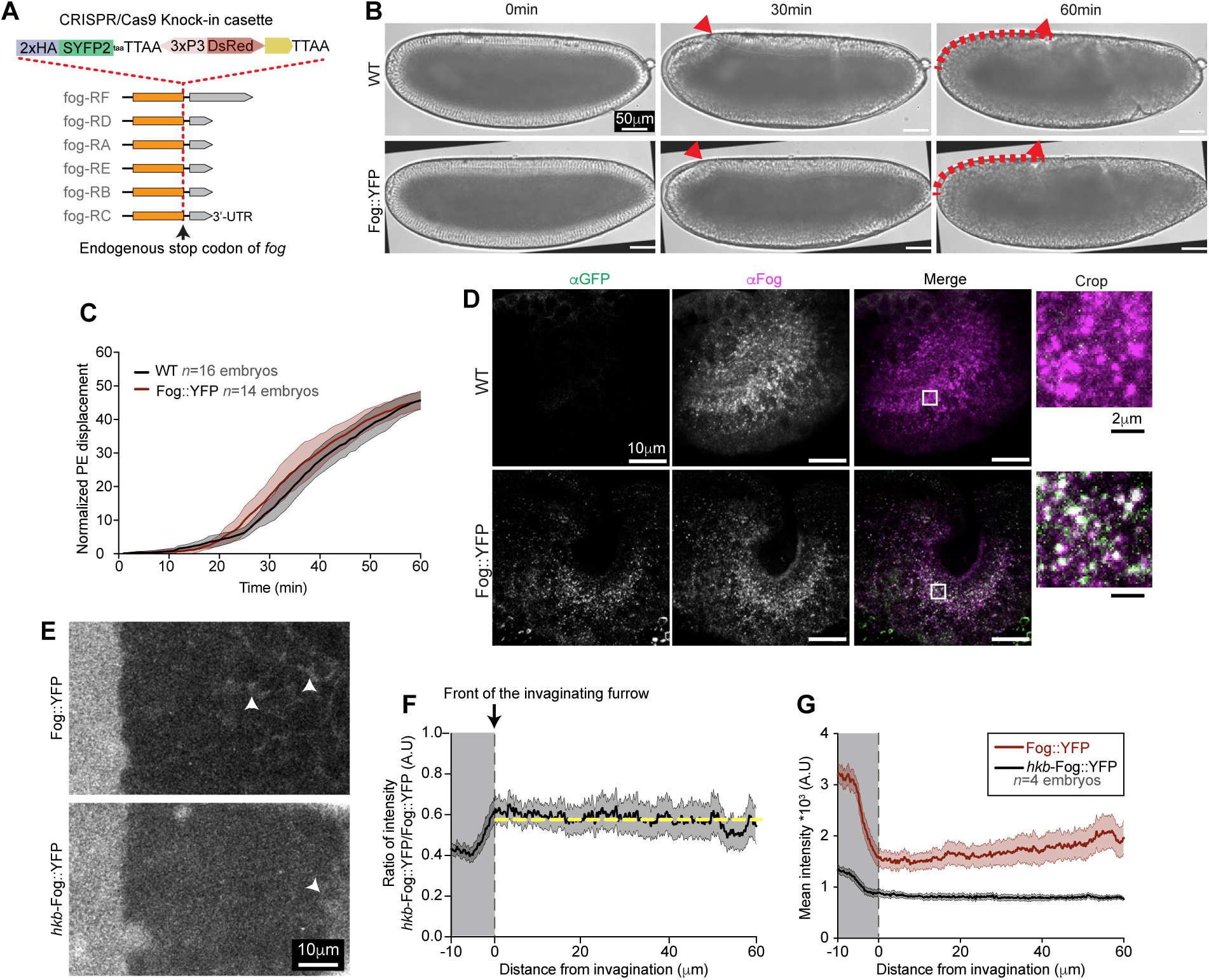
Endogenously tagged Fog::YFP shows similar localization to WT Fog. (**A**): CRISPR/Cas9-based strategy for genome engineering of endogenously tagged Fog with a SYFP2 (see methods). (**B**): Stills from brightfield time-lapses of WT and Fog::YFP embryos. The red arrowheads mark the dorsal edge of the endoderm invagination and red dashed lines indicate the displacement of the posterior endoderm at the indicated times. (**C**): Quantification of the normalized posterior endoderm (PE) displacement over time in WT and Fog::YFP embryos. Time 0 is the time when the cellularization front reaches the basal side of nuclei on the dorsal side of the embryo. *n* =16 and 14 embryos for WT and Fog::YFP respectively. (**D**): Representative sagittal sections of IHC of Fog and GFP in WT (top) and Fog::YFP (bottom) embryos at wave propagation stage during endoderm morphogenesis. On the right, zoomed merge views of the indicated white box on the left to show the colocalization of Fog (magenta) and GFP (green). (**E**): Extracellular distribution of endogenous (Fog::YFP) and zygotically produced Fog::YFP (*hkb-fog::SYFP2,* referred to as *hkb*-Fog::YFP) during wave propagation. White arrows indicate Fog present in the extracellular space anterior to the invaginating furrow. (**F**): Spatial profile of the ratio of mean intensities between Fog::YFP and hkb-Fog::YFP. The yellow dashed line indicates a constant ratio. The moving front of the invagination is at 0μm. (**G**): The spatial mean intensity profiles of indicated conditions. In **F** and **G** *n*=4 embryos Statistics: Data in **C** is mean±s.d and data in **F** is mean±SEM. Scale bars, 50μm (**B**), 10 μm (**D** and **E**) and 2 μm (**D**, crop).

**Figure S5.**
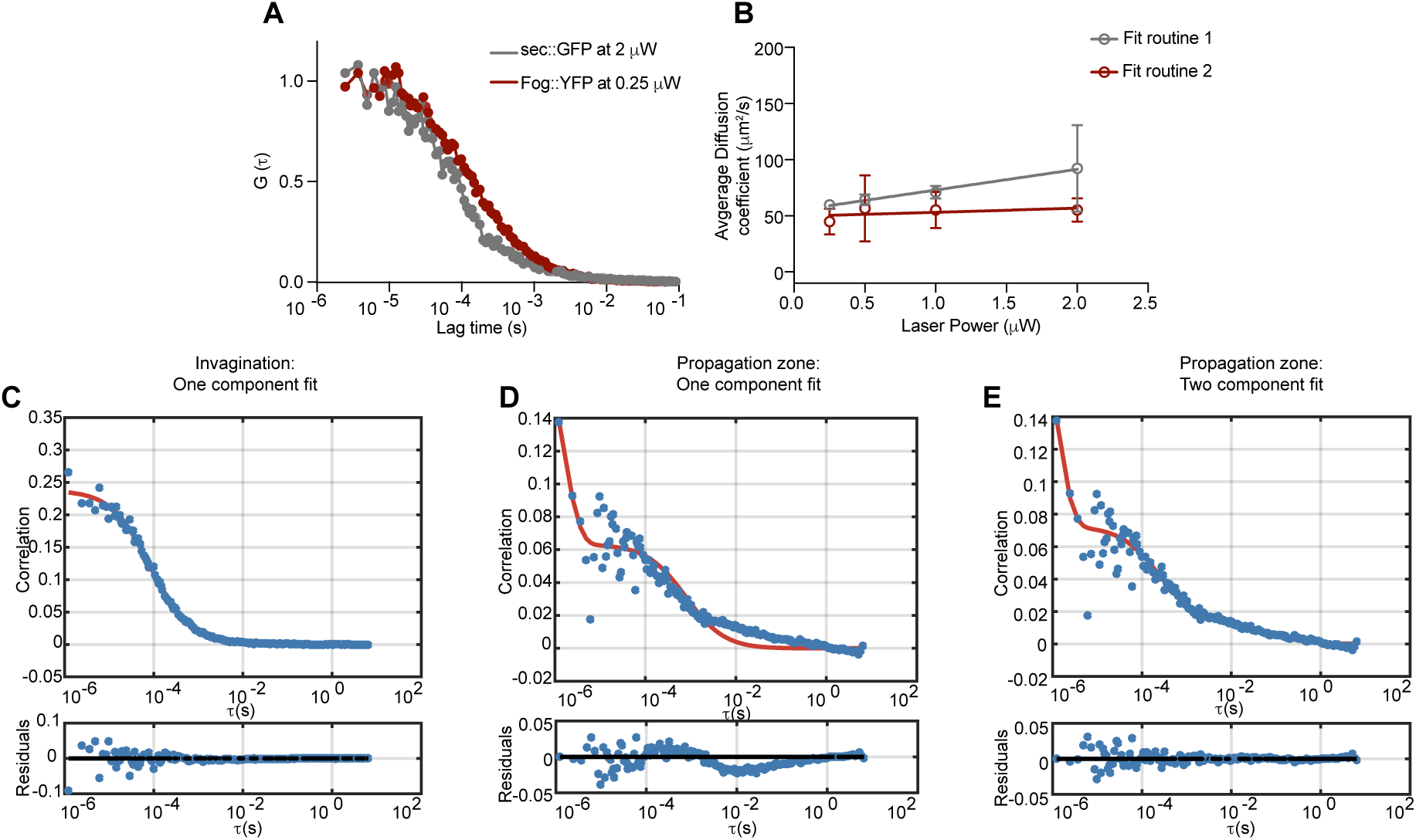
FCS curve-fitting. (**A**): Autocorrelation function (G(τ)) obtained by averaging multiple FCS measurements for Fog::YFP (*n* =9) and sec::GFP (*n* =8) in the invagination, acquired at comparable effective excitation powers. Fitting these average ACFs was used to extract diffusion coefficients shown in Fig. 4H. Fitting of the individual ACFs was used to extract the D values shown in Fig. 4G. (**B**): Diffusion coefficients for Fog::YFP obtained by fitting two different fit routines to ACFs acquired at different laser powers (see methods). Fit routine 2 mitigates the bias towards larger D values caused by photophysical artifacts. (**C-D**): Representative ACFs (blue circles) from FCS measurements of Fog::YFP in the invagination (**C**) or in the propagation zone (**D**) fitted with a one-component diffusion model (red line in the top graph). On the bottom the fit residuals are shown. (**E**): Representative ACF (blue circles) from the same FCS measurement in the propagation zone fitted with a two-component diffusion model (red line in the top graph). On the bottom the residuals are shown.

**Figure S6.**
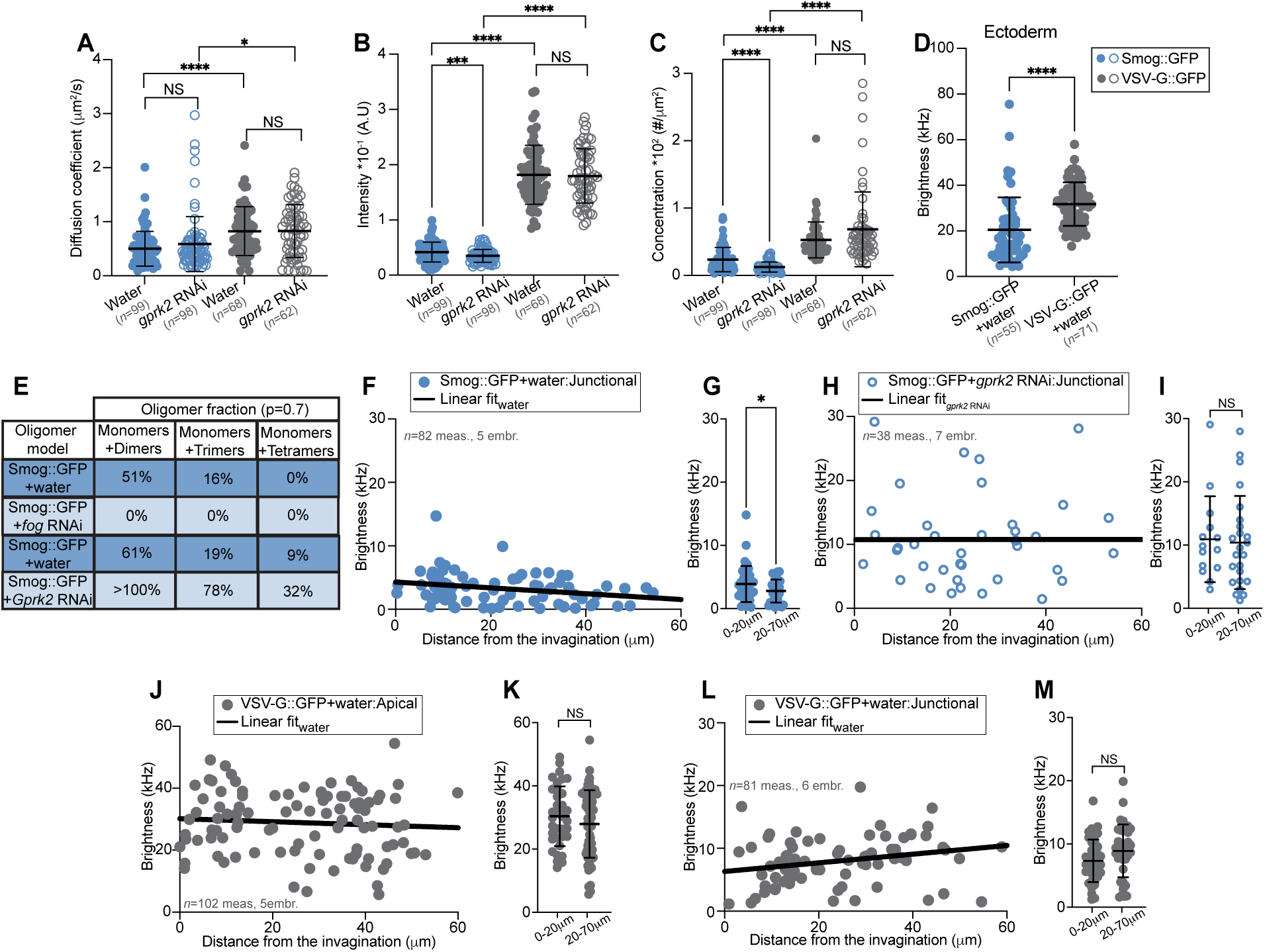
Slowly diffusing Fog::YFP and Smog::GFP homoclusters form a gradient at the junctions in the PZ. (**A-C**): Measurements of Smog::GFP (blue) and VSV-G::GFP (grey) diffusion coefficients (**A**), average intensity (**B**) and molecular concentration (**C**) at the apical membrane in cells of the PZ in water (control) and *gprk2* RNAi injected embryos during wave propagation. *n=*99 measurements from 14 embryos (Smog::GFP + water), 98 measurements from 20 embryos (Smog::GFP + *gprk2*RNAi), 68 measurements from 5 embryos (VSV-G::GFP + water) and 62 measurements from 8 embryos (VSV-G::GFP + *gprk2* RNAi). In **A**, NS is *P*=0.5202 (Smog::GFP + water vs Smog::GFP + *gprk2* RNAi), *P*=0.9885 (VSV-G::GFP + water vs VSV-G::GFP + *gprk2* RNAi), \**P*=0.0377, \*\*\*\**P*<0.0001 unpaired *t*-test with Welch’s correction. In **B**, NS is *P*=0.8087, \*\*\**P*=0.0002, \*\*\*\**P*<0.0001 unpaired *t*-test with Welch’s correction. In **C,** NS is *P*=0.0930, \*\*\*\**P*<0.0001 unpaired *t*-test with Welch’s correction. (**D**): Molecular brightness of the indicated molecules in the lateral ectoderm during germband elongation in indicated conditions. *n=*55 measurements from 11 embryos (Smog::GFP + water) and 71 measurements from 6 embryos (VSV-G::GFP + water). \*\*\*\**P*<0.0001 unpaired *t*-test with Welch’s correction. (**E**): Estimation of the fraction of monomers and higher-order oligomers (see methods) of Smog::GFP at the apical surface of cells in the PZ during wave propagation in indicated conditions, calculated under the assumption of limited GFP maturation and presence of dark fluorophore states (i.e., p<1, here p=0.7 was assumed). ‘p’ is the probability of each GFP label being fluorescent (see methods). (**F**) and (**H**): Molecular brightness of Smog::GFP measured at junctions at different distances from the invaginating furrow in the PZ during wave propagation in water (**F**, *n=* 82 measurements from 5 embryos) and *gprk2* RNAi (**H**, *n=*38 measurements from 7 embryos) injected embryos. (**G**) and (**I**): Junctional Smog::GFP molecular brightness for the indicated bins of distance in water (**G**, *n=*82 measurements from 5 embryos, \**P*=0.0455) and *gprk2* RNAi (**I**, *n=*38 measurements from 7 embryos, *P*=0.8241) injected embryos. (**J**) and (**L**): Molecular brightness of VSV-G::GFP measured at the apical surface (**J**, *n=*102 measurements from 5 embryos) and junctions (**L**, *n=* 81 measurements from 6 embryos) at different distances from the invaginating furrow in the PZ during wave propagation in WT embryos. (**K**) and (**M**): Apical (**K**) and junctional (**M**) VSV-G::GFP molecular brightness for the indicated bins of distance in WT embryos. In **K**, *n=*102 measurements from 5 embryos, *P*=0.2359. In **M,** *n=*81 measurements from 6 embryos, *P*=0.0735. In **F**, **H**, **J** and **L** the solid black line represents a simple linear regression fitting. Statistics: Data are mean±s.d.

**Figure S7.**
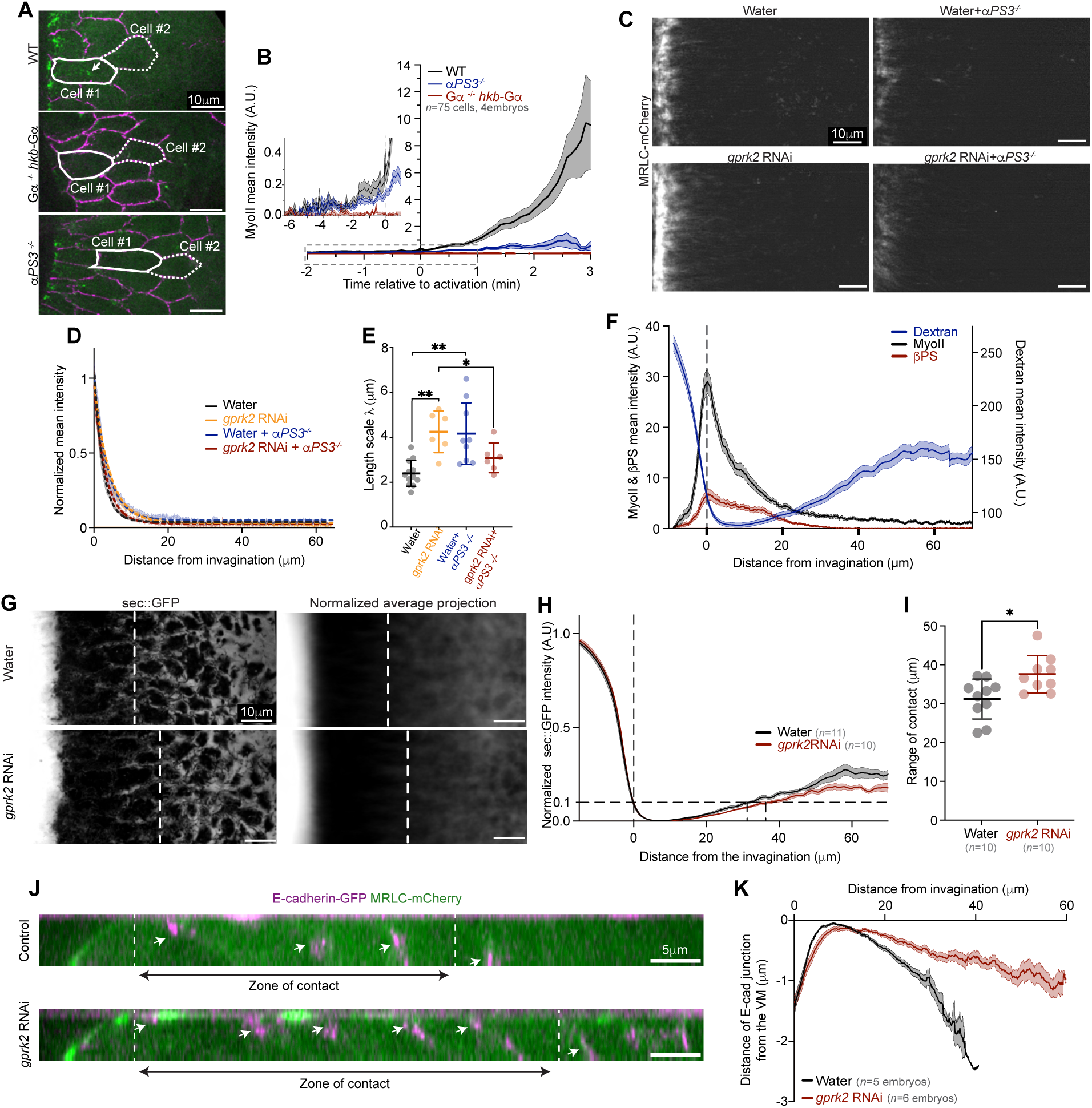
Integrins amplify MyoII activation over a long spatial range and the contact with the VM is expanded in *gprk2* RNAi. (**A**): Stills from time-lapses of MyoII in a WT (top), a *Gα*^−/−^ *hkb*-Gα (middle) and an integrin mutant (*αPS3^−/−^*, bottom) embryo. (**B**): MyoII mean intensity over time in cells of the PZ in the indicated conditions. Time 0 for each cell is the time when cell area is maximum (see methods). The inset is a zoomed view of the boxed region. *n=*75 cells across 3 embryos each. (**C**): Temporal average projections of MyoII in the indicated conditions. (**D**): Normalized spatial profiles of the MyoII mean intensity in indicated conditions (same data as in Fig. 6B). (**E**): Measurements of the length scale of MyoII in indicated conditions. In **C**, **D** and **E** *n*=12 embryos (Water), 8 embryos (*gprk2* RNAi), 9 embryos (water + *αPS3^−/−^*) and 6 embryos (*gprk2* RNAi + *αPS3^−/−^*). \*\**P*=0.0029 (Water vs *gprk2* RNAi), \*\**P*=0.0046 (Water vs *αPS3^−/−^*) and \**P*=0.0366, unpaired *t*-test with Welch’s correction. (**F**): Average spatial profiles of MyoII, βPS and dextran mean intensity in the PZ of WT embryos. *n*=9 embryos each. (**G**): Left: Stills form timelapses of secreted GFP (sec::GFP) in water (top) and *gprk2* RNAi injected (bottom) embryos. Right: Normalized temporal average projections in the referential of the moving invagination (see methods) of sec::GFP in the indicated conditions. The dashed white line represents the zone where the tissue is in contact with the VM. *n*=11 embryos (Water), 10 embryos (*gprk2* RNAi). (**H**): Normalized spatial profiles of the sec::GFP mean intensity in the PZ in the indicated conditions. 0µm indicates the position of the invagination. *n*=11 embryos (Water), 10 embryos (*gprk2* RNAi). The dashed horizontal line indicates the activation threshold estimated as described in the methods. (**I**): Measurements of the range of tissue contact with the VM in front of the invaginating furrow estimated from the exclusion of dextran (see methods). *n*=11 embryos (Water), 10 embryos (*gprk2* RNAi). \**P*=0.0315, unpaired *t*-test with Welch’s correction. (**J**): Side views of E-cad and MyoII in water and *gprk2* RNAi injected embryos. The white arrows indicate the position of E-cad junctions. (**K**): Spatial profile of the contact to the VM measured as the distance of E-cadherin junctions from the VM. *n*= 5 embryos (Water), 6 embryos (*gprk2* RNAi). Statistics: Data in **E** and **I** are mean±s.d. Data in **B**, **F**, **H** and **K** are mean±e.e.m. Scale bar, 10 μm.

## Movie Legends

**Movie S1**

Tissue-level MyoII dynamics during endoderm morphogenesis. Time-lapse of the morphogenesis of the posterior endoderm in a WT (top) and a *Gα* ^−/−^ *hkb-*G*⍺* (bottom) embryo. E-cadherin-GFP is in magenta and MRLC-mCherry in green. Scale bar 15µm. N= 5 embryos.

**Movie S2**

Cell level MyoII dynamics in the primordium region. Close-up views from a time-lapse of MyoII recruitment in a cell in the primordium region in a WT (left) and a *Gα*^−/−^ *hkb-*Gα (right) embryo. E-cadherin-GFP is in magenta and MRLC-mCherry in green. The yellow contour marks a tracked cell over time. Scale bar 5μm. N= 5 embryos.

**Movie S3**

Gastrulation in *Gα*^−/−^*hkb-*Gα embryos. Bright-field time-lapse of gastrulation in a WT (top) and a *Gα^−/−^hkb-* Gα (bottom) embryo. Scale bar 50μm. N= 25 embryos.

**Movie S4**

MyoII activation in the propagation zone. High-resolution time-lapse of cells in the propagation zone in a WT (top) and *Gα*^−/−^ *hkb-*Gα (bottom) embryo. E-cadherin-GFP is in magenta and MRLC-mCherry in green. Scale bar 10μm. N= 5 embryos.

**Movie S5**

Tissue-level MyoII dynamics in embryos injected with *fog* RNAi and expressing different versions of fog under the *hkb* promoter. On the top a time-lapse of an embryo expressing an RNAi-sensitive version of *fog* (*hkb-fog_sen_*). In the middle an embryo expressing an RNAi-resistant version of *fog* (*hkb-fog_res_*). On the bottom an embryo expressing an RNAi-resistant version of *fog* tethered to the membrane in the primordium region (*hkb-fog_VSV-G_*). E-cadherin-GFP is in magenta and MRLC-mCherry in green. Scale bar 15μm. N= 5 embryos.

**Movie S6**

Gastrulation visualized by bright field imaging in *fog* RNAi embryos expressing different versions of fog under the *hkb* promoter. On the top an embryo expressing an RNAi-sensitive version of *fog* (*hkb-fog_sen_*). In the middle an embryo expressing an RNAi-resistant version of *fog* (*hkb-fog_res_*). On the bottom an embryo expressing an RNAi-resistant version of *fog* tethered to the membrane in the primordium region (*hkb-fog_VSV-G_*). Scale bar 50μm. N= 25 embryos.

**Movie S7**

Cell level MyoII dynamics in the primordium region in embryos injected with *fog* RNAi and expressing different versions of fog under the *hkb* promoter. Close-up views from a time-lapse of MyoII recruitment in a cell in the primordium region in a *hkb-fog_res_* (left) and *hkb-fog_VSV-G_* (right) embryo. E-cadherin-GFP is in magenta and MRLC-mCherry in green. The yellow contour marks a tracked cell over time. Scale bar 5μm. N= 5 embryos.

**Movie S8**

MyoII activation in the propagation zone in embryos injected with water or *fog* RNAi and expressing different versions of fog under the *hkb* promoter. High-resolution time-lapse of cells in the propagation zone in a *hkb-fog_res_* injected with water (top), *hkb-fog_res_* injected with *fog* RNAi (middle) and *hkb-fog_VSV-G_* injected with *fog* RNAi (bottom) embryo. E-cadherin-GFP is in magenta and MRLC-mCherry in green. Scale bar 10μm. N= 5 embryos.

**Movie S9**

MyoII pattern in embryos where diffusible or membrane-tethered Fog are expressed ubiquitously. Time-lapse of MyoII activation during endoderm morphogenesis in embryos homogenously expressing a diffusible (UAS-*fog*^12^, top) or plasma membrane-tethered version (UAS-*fog_VSV-G_*, bottom) of Fog. Scale bar 15μm. N= 5 embryos.

**Movie S10**

MyoII dynamics of the propagation zone upon increased Fog production in the endoderm primordium. High-resolution time-lapse of cells in the propagation zone in a WT (top) and an embryo where *fog* expression is increased in the primordium (*hkb*-*fog*, bottom). E-cadherin-GFP is in magenta and MRLC-mCherry in green. Scale bar 10μm. N= 4 WT embryos and 9 *hkb*-Fog embryos.

**Movie S11**

MyoII dynamics in the propagation zone upon reduced Fog endocytosis. High-resolution time-lapse of cells in the propagation zone in WT (top) and embryos uniformly expressing shRNAs targeting GPRK2 (*gprk2* RNAi, bottom). E-cadherin-GFP is in magenta and MRLC-mCherry in green. Scale bar 10μm. N= 5 WT embryos and 9 *gprk2 RNAi* embryos.

**Movie S12**

Gastrulation in embryos with endogenously tagged Fog. Bright-field time-lapse of a WT (top) and a Fog::YFP (bottom) embryo. Scale bar 50μm. N= 16 WT and 14 Fog::YFP embryos.

**Movie S13**

Fog protein distribution during gastrulation. Confocal time lapse in the midsagittal plane of an embryo expressing endogenously tagged Fog with SYFP2 (Fog::YFP) imaged on a two-photon microscope. Scale bar 50μm. N= 5 embryos.

**Movie S14**

Fog protein distribution during endoderm morphogenesis. Confocal time-lapse in the posterior endoderm of Fog::YFP embryo. Scale bar 20μm. N= 5 embryos.

**Movie S15**

Secreted GFP protein distribution during endoderm morphogenesis. Confocal time-lapse in the posterior endoderm of an embryo uniformly expressing a secreted form of GFP (sec::GFP). Scale bar 20μm. N= 5 embryos.

**Movie S16**

Distribution of zygotically produced Fog::YFP compared to the distribution of the entire pool (maternal + zygotic). Confocal time-lapse of endogenously tagged Fog (Fog::YFP, top) and zygotically produced *fog* expressed under the *hkb* promoter (*hkb-*Fog::YFP, bottom). Scale bar 10μm. N= 5 embryos.

**Movie S17**

MyoII dynamics in the propagation zone upon reduced Fog endocytosis in integrin mutants. High-resolution time-lapse of the propagation zone in water-injected embryos (water, top left), water-injected integrin mutants (water+*αPS3*^−/−^, top right), *gprk2* RNAi-injected embryos (*gprk2* RNAi, bottom left) and *gprk2* RNAi-injected integrin mutants (*gprk2*RNAi +*αPS3^−/−^*, bottom right). E-cadherin-GFP is in magenta and MRLC-mCherry in green. Scale bar 10μm. N= 5 embryos.

**Movie S18**

Integrin and MyoII dynamics during wave propagation in WT and *gprk2* RNAi-injected embryos. High-resolution time-lapse of βPS (left) and MyoII (right) in embryos injected with water (top) or *gprk2* RNAi (bottom). Scale bar 10μm. N= 6 water-injected and 11 *gprk2* RNAi-injected embryos.

**Movie S19**

The zone of contact with the vitelline membrane is expanded in *gprk2* RNAi-injected embryos. High-resolution time-lapse of apical planes just below the vitelline membrane in embryos uniformly expressing sec::GFP injected with water (top) or *gprk2* RNAi (bottom). Scale bar 10μm. N= 11 water-injected and 10 *gprk2* RNAi-injected embryos.

## Materials and methods

### Fly strains and genetics

The following mutant alleles and insertions were used:

*cta^RC10^*(gift from Leptin lab), *pUASt-fog^1^* (*5*), *pUASp-sec::eGFP* (*46*), *hkb-*fog (*32, 33*), *pUAS-gprk2shRNA* (Bloomington stock #35326), *scb*^KO^ (*30*), *ubi-VSV-G::GFP* (*37*), *sqh-smogC::GFP* (long ORF of the RC isoform of Smog, chromosome 3 (*37*) were described earlier. *hkb-Gα*, *hkb-fog_res_*, *pUASt-fog_VSV-G_*, *hkb-fog_VSV-G_*, and *fog::SYFP2^KIN^* were generated in this study (see below).

Live-cell imaging of MRLC, *spaghetti squash* (*sqh*, Genebank ID: AY122159) in *Drosophila* was carried out using a *sqh-Sqh::mCherry* transgene inserted either on chromosome 2 (at the VK18 site, located at 53B2 (*6*) or a previously generated insertion by Adam Martin (*22*)) or on chromosome 3 (VK27 site located at 89E11 (*47*)). Live imaging of E-cadherin (*shg* in *Drosophila*, FlyBase ID:FBgn0003391) and of the main beta integrin βPS (*myospheroid (mys)*, in Drosophila FlyBase ID: FBgn0004657) was carried out with EGFP knock-in alleles at the locus generated by homologous recombination, respectively *E-cad::EGFP^KIN^* (*48*) and *mys::EGFP^KIN^* (*49*).

For the expression of Fog in stripes (Fig. S3B,C), females ;;*wg-Gal4* were crossed to males ;;*pUASt-fog^12^* or ;;UAS-fog_VSV-G_ and the F1 progeny (embryos) were analyzed. For the homogeneous overexpression of Fog (Fig. S3E), *females* ;*67-Gal4*,*E-cad::EGFP^Kin^*,*sqh-Sqh::mCherry* were crossed with males ;*pUASt-fog^12^; or* ;;*pUASt-fog_VSV-G_* and the F1 progeny was analysed. The presence of the UAS transgene was assessed based on the phenotype. 67-Gal4 (mat α4-GAL-VP16) is ubiquitous and maternally supplied.

For the expression of shRNA against GPCR kinase-2 (*gprk2*, FlyBase ID: FBgn0261988), F1 progeny was analzyed from females ;*67-Gal4*,*E-cad::EGFPKin,sqh-Sqh::Cherry/+*; *pUAS-gprk2shRNA*/+ crossed to males ;;*pUAS-Gprk2shRNA*.

For homogenous expression of secreted GFP (*sec::GFP*), females ;*67-Gal4*/+;*UAS-sec::GFP/+* were crossed to males *;;UAS-sec::GFP* and F1 progeny was analzyed.

All fly constructs and genetics are listed below.

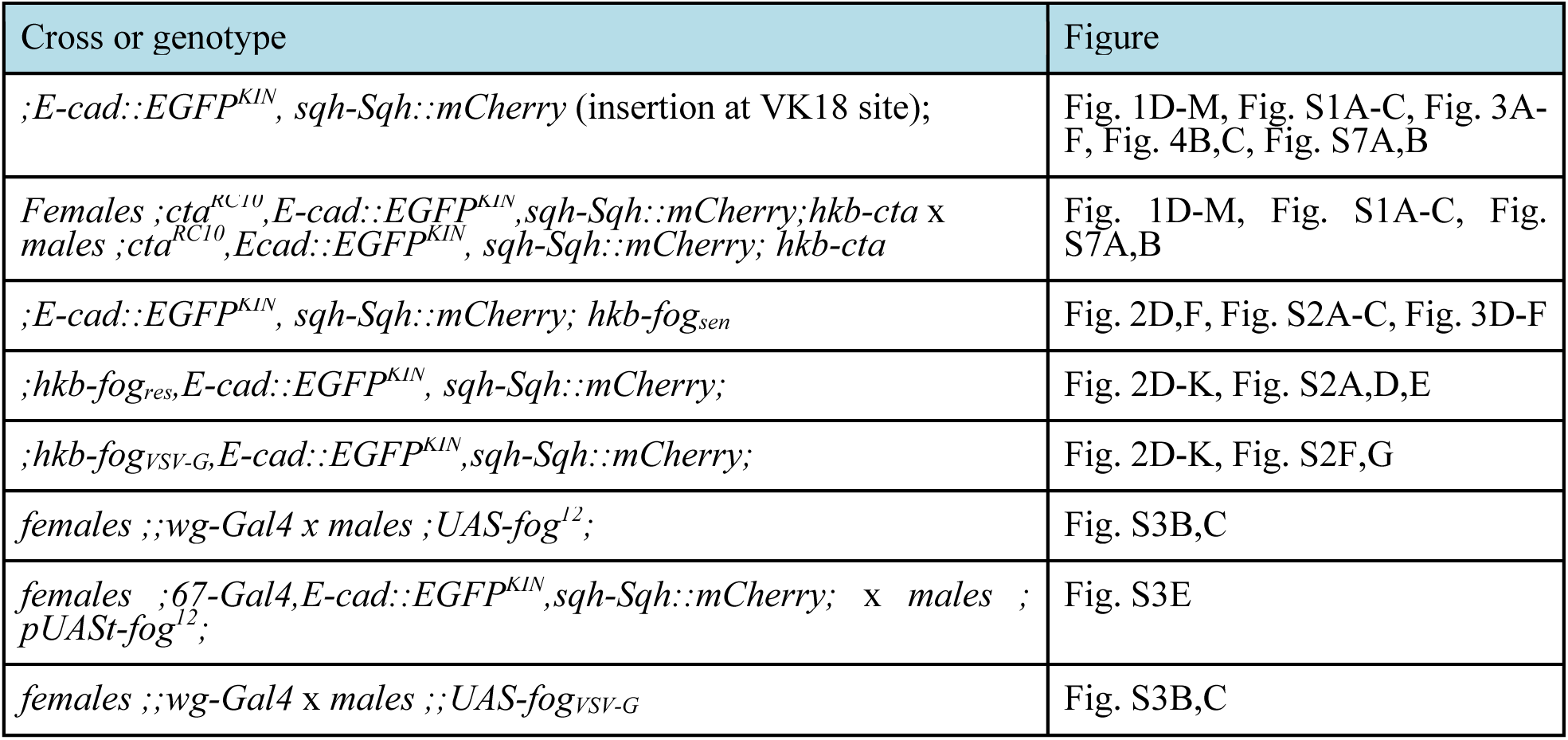
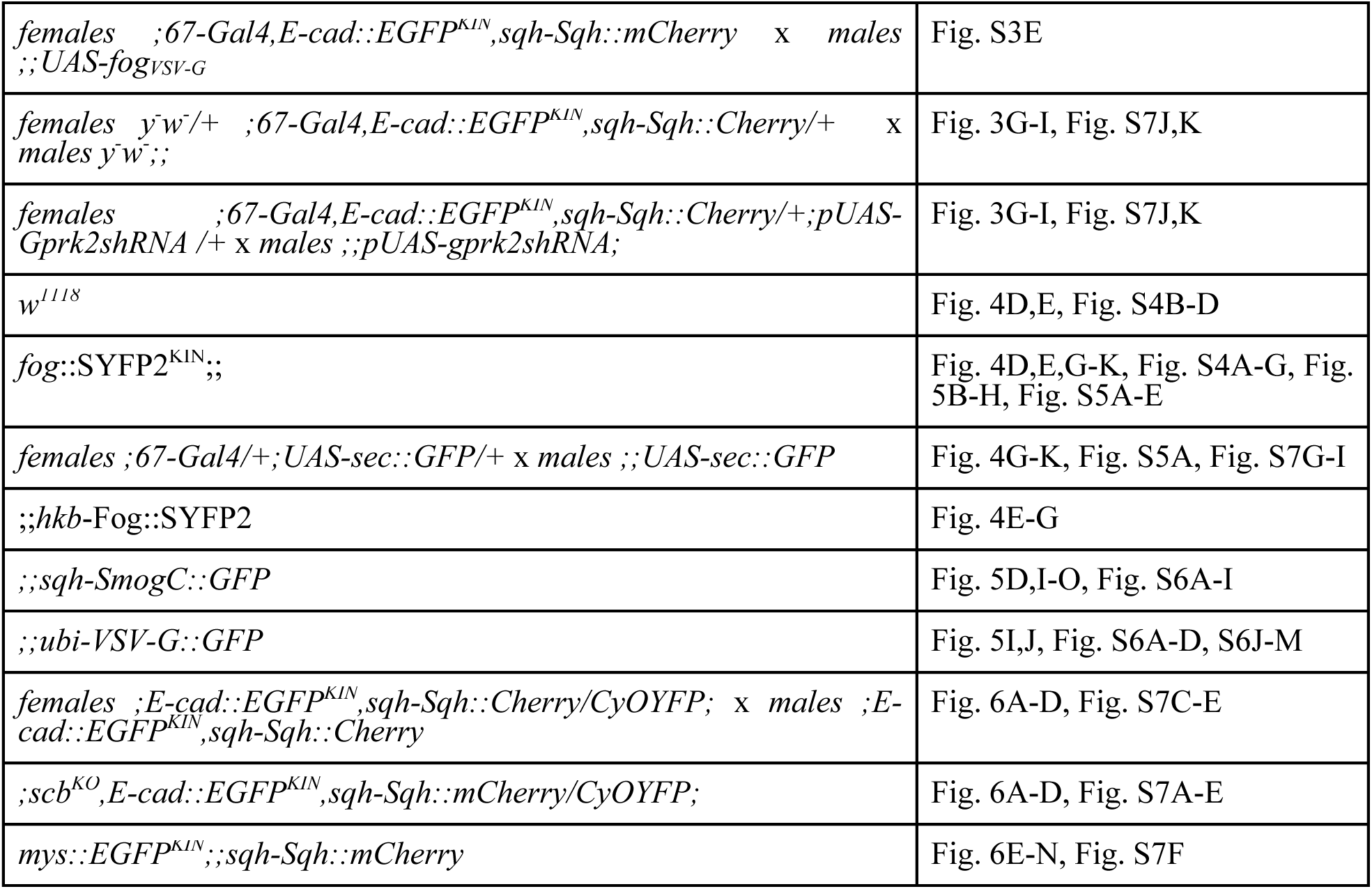

### Constructs and transgenesis

#### *hkb*-Gα

We re-built the plasmid containing *hkb-fog* from Ref. (*33*). The sequence from nucleotide −2498bp upstream the ATG to the ATG itself of the *hkb* gene was cloned upstream an hsp70 basal promoter and the complete fog-RB mRNA (NM_001038771.3) into a pCasper5 transformation vector containing an attB site for PhiC31-mediated transgenesis. The pCasper5-attB backbone was purified after EcoRI digest of pBR14 (Addgene 52522, a pCasper5-attB_Cas9 plasmid from Klaus Foerstemann (*50*)). This was called the C5-hkb-Fog plasmid. The *fog* ORF of in the C5-hkb-Fog plasmid was replaced with the Gα ORF (cta, NM_001273767.1) for building *hkb*-Gα.

#### hkb-fog_res_

To build a *fog* mRNA resistant to the dsRNA (*fog_re_*_s_) under the control of the *hkb* promoter, the sequence of fog-RB mRNA targeted by the injected dsRNAs (861bp: from 24bp before the ATG to 834bp after the ATG) in the C5-hkb-Fog plasmid was modified every 4 to 8 bp using codon redundancy to generate the same aminoacidic sequence (see Fig. S2A, detailed sequence available in Table S1).

#### hkb-fog_VSV-G_

Membrane-tethered Fog was built using the *fog* ORF (U03717.1) fused in C-terminal with a truncated version of type-I transmembrane protein VSV-G (vesicular stomatitis virus-G protein, Genebank NP 041715, amino acids E422-R508, ACK77584). Specifically, the extracellular amino acids 1 to 421 of the VSV-G protein were deleted and 2xHA tags were inserted between the *fog* ORF and the truncated VSV-G ORF surrounded by two SGGGGS flexible linkers. The fusion construct was cloned in the pCasper5 plasmid backbone. *hkb-fog_VSV-G_* was inserted into the attP2 landing site (Chr3, 68A4) and *hkb-fog_res_* into the attP40 landing site (Chr2, 25C6).

#### scab^ko^ allele

a KO-attP founder line was generated by CRISPR/Cas9 gene editing as previously described (*30*). Briefly, the entire ORF and a part of the UTRs were deleted (from −281 nucleotide to +8,670 nucleotide from ATG of scab-RB, Id=FBtr0087369) and replaced by an attP-3xP3-RFP selection marker cassette containing: forward attP (49bp) and a floxed 3xP3-RFP eye selection marker which was flipped out by Cre recombinase in a second step. The deletion was verified by genomic PCR and Sanger sequencing. This “KO-attP founder” allele is a null allele of *scab*. Flies are not homozygous viable and are maintained over CyOdYFP balancer.

#### *fog*::SYFP2^KIN^

We used SYFP2 (*51*) to generate endogenously tagged Fog. SYFP2 was used for tagging because it is reported to be a very rapidly maturing monomer (*51*). Endogenous *fog* tagged to SYFP2 (*fog*::SYFP2^KIN^) was generated by CRISPR/Cas9 gene editing (Wellgenetics Fly Genome Editing Service, Taipei, Taiwan), using a donor vector containing in sequence (Fig. S4A): 1) the 5’-fog homology arm, 2) a 2xHA tags flanked by two SGGGGS flexible linkers, 3) the SYFP2 inserted before the TAA stop codon of the *fog* gene, 4) a PiggyBac DsRed eye marker screening cassette, composed of 3xP3-selection cassette (*52*) flanked by the 5’ and 3’ PiggyBac terminal repeats, and cloned just after the TAA stop codon and 5) the 3’-fog homology arm. Following eye colour selection to identify successful gene editing, the 3xP3-DsRed cassette was flipped out by PiggyBac Transposase generating a *fog* allele (*fog*::SYFP2^KIN^) encoding a Fog::YFP fusion protein at the endogenous locus. The *fog*::SYFP2^KIN^ allele is homozygous viable and shows no defects in germband extension (Fig. 4B,C). Correct gene editing was verified by genomic PCR and Sanger sequencing.

FASTA sequences of all plasmids are available upon request.

### RNA interference

The dsRNA probe against *fog* was prepared as previously described (*6, 34*). The probe is 861 bp long (from 24bp upstream to the ATG to 834bp downstream) and targets nucleotides 1546-2406 of the *fog* (CG9559) mRNA (Genebank, NM_078714). dsRNA probes were prepared and injected at a final concentration of 5-30 μM in embryos less than 1h old as previously described (*34, 53*). The sequence of the primers used to generate the dsRNA probes against *fog* is shown below (underline is the annealing sequence and the remaining is the T7 promoter).

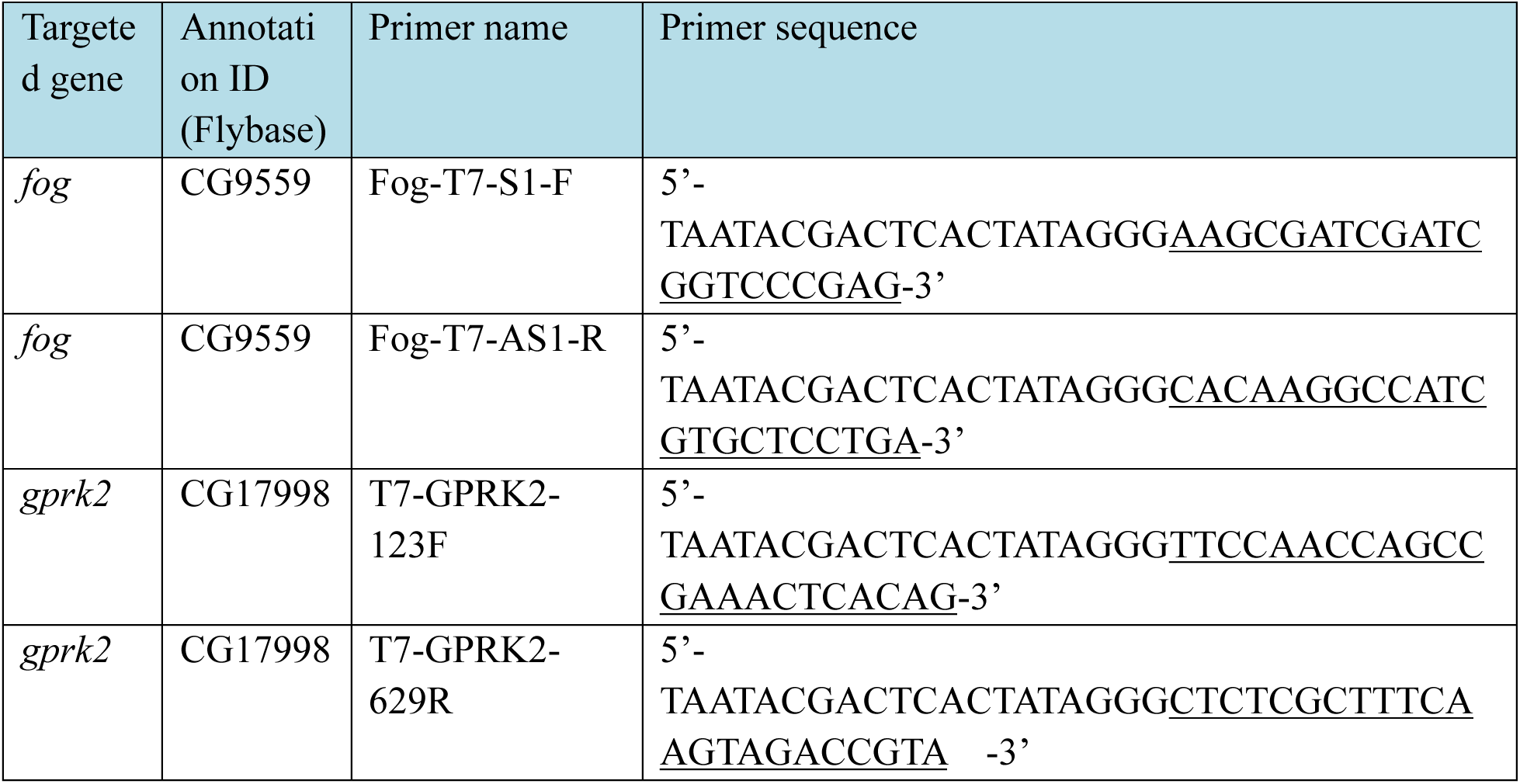

The dsRNA against *gprk2* was prepared as previously described (*37*). The dsRNA probe against *gprk2* (CG17998) is 529 bp long and targets nucleotides 659-1,187 located in 5’-UTR of the gprk2-RA transcript (GenBank ID:NM_057519). A 5μM dilution of the probe was injected into embryos less than 1h old. The sequence of the primers used to generate the dsRNA probes against *gprk2* is shown below in Table 2 (underline is the annealing sequence and the remaining is the T7 promoter).

Dextran injections (Fig. 6E, Fig. S7F) were performed as previously described (*6*), using dextran 647, 10,000MW, anionic, fixable from Thermo Fischer Scientific (D22914) at a concentration of 1 mg/mL. Embryos were injected in the perivitelline space at stage 6-7 directly on the microscope and imaged few minutes later to ensure dextran equilibration in the perivitelline fluid.

All the injections were performed at 50-80% embryo length from the posterior and on lateral side in all experiments.

### Sample preparation

Flies were kept in cages at 25°C except for experiments using the UAS-GAL4 system (*wg-Gal4* and *67-Gal4*) in which they were kept at 18°C. Embryos were prepared as previously described (*54*). Briefly, they were collected using apple cider plates smeared with yeast paste and transferred to a mesh basket, rinsed with water, dechorionated with 2.6% bleach for 90 seconds, and then rinsed copiously with water before transferring them back to clean agar. For live imaging, embryos were staged and selected (early stages of cellularization) under a dissection microscope and were aligned with the dorsal side facing up. They were then transferred to a glass coverslip coated with homemade glue. A drop of Halocarbon 200 Oil (Polysciences) was placed on the embryos to avoid drying during imaging.

### Live imaging

Live imaging of embryos was performed at stage 7 in the dorsal-posterior region for 12-50 min at room temperature (21-22°C), depending on the experiment. Dual colour time-lapse imaging for EGFP and mCherry was performed using simultaneous acquisition on a spinning disc confocal (CSU-X1, Yokogawa) Nikon Eclipse Ti inverted microscope equipped with two cameras (Rolera EM-C^2^, Q-Imaging or Kinetix22, Photometrics) distributed by Roper and using a 40X/1.25NA Apo water-immersion or a 100X/1.45NA Plan Apo oil-immersion objectives from Nikon, depending on the experiment. Live imaging was performed by simultaneously exciting with 491-nm (EGFP) and 561-nm (mCherry) lasers, using a dichroic mirror to collect emission signals on two cameras. For live imaging with 40X magnification, z-series of 33 planes spanning (spacing 0.8μm) spanning about 26.4μm from the apex of cells in the PZ was acquired with a frame rate of 1 stack every 20 s. For live imaging of E-cad and MyoII with 100X magnification, z-series of 10 planes spanning 5μm (spacing 0.5μm) was acquired with a frame rate of 1 stack every 5 s. Live imaging of sec::GFP in Fig. 4H was performed with 100X magnification, z-series of 10 planes spanning 5μm (spacing 0.5 μm) below the vitelline membrane and with a frame rate of 1 stack every 5s. Live imaging of βPS and MyoII was performed with 100X magnification. z-series of 2 planes spanning 1μm (spacing 0.5 μm) just below the vitelline membrane were acquired with a frame rate of 1 stack every 4 s. Similar settings were used to image sec::GFP in Fig. S7G.

Live imaging of endogenous Fog::YFP and *hkb*-Fog::YFP was performed with a Zeiss 880 confocal microscope using a 40X/1.2NA water immersion or 63X/1.4 NA oil objective. GaAsP hybrid detectors were utilized with a 514-nm excitation laser. Z-stacks of 4-5 planes (spacing 0.5 μm) spanning about 2-2.5μm just below the vitelline membrane was acquired every 15-90 s for 15-45 min.

Imaging conditions (line averaging, camera exposure time, laser power, etc.) were optimized and kept constant during the experiment performed at room temperature (22 °C). Laser powers were measured with a power meter at the back aperture of the objectives for every imaging session.

For brightfield imaging in Fig. S1B, Fig. S2B,D,F and Fig. S4B, embryos were prepared and immobilized for imaging as described above. Bright-field time-lapse images were collected on a Zeiss Axiovert 200M inverted microscope using a 20X/0.75NA objective and a programmable motorized stage to record different positions over time (Mark&Find module from Zeiss). The system was controlled by AxioVision software (Zeiss). Time-lapse data in WT or injected embryos were performed over 300 mins with a frame rate of 1 image every 1min.

### Antibody staining

Embryos were fixed and permeabilized with 3.7% formaldehyde for 20 min, the vitelline membrane was removed by shaking in heptane/methanol and then embryos were stained according to standard procedures (*55*). The Fog protein was detected with a rabbit antibody (1:1000, a gift from N.Fuse (*35*), *sqh-MRLC::mCherry* was detected with a rat antibody (1:1000, anti-RFP, Chromotek 5f8) and Fog::YFP was detected with a chicken antibody (1:1000, anti-GFP, Aveslabs, GFP-1020). The Fog antibody was pre-adsorbed by incubation overnight at 4℃ with fixed *y^−^w^−^* embryos less than 1hr old. Patched was detected with a mouse antibody (1:100, Drosophila Ptc (apa1) from DSHB, AB_528441). Secondary antibodies used were donkey anti-rabbit Alexa Fluor 568 (1:500, Thermo Fischer Scientific, A10042), donkey anti-rat Alexa Fluor 568 (1:500, Thermo Fischer Scientific, A78946), donkey anti-chicken Alexa Fluor 488 IgY (1:500, Thermo Fischer Scientific, 703 545 155), donkey anti-mouse Alexa Fluor 647 (1:500, Thermo Fischer Scientific, A32787). Stained embryos were mounted in Aqua-Polymount (Polysciences) and imaged with a Zeiss 880 confocal microscope using a C-Apochromat 63X/1.4 NA oil objective. Image stacks with spacing of 0.5 μm were collected and maximum projections of 3–7 planes were analysed.

### Image processing, segmentation and cell tracking

All image manipulation and processing were performed using ImageJ (version 2.140/1.54f). For 40X image stacks, image projections of selected z-planes around the cell apex were performed as previously described (*6*) using a custom procedure exploiting the Stack Focuser plugin (M. Umorin). The cells were then tracked either manually (by manually drawing ROIs) or automatically (Tissue Analyzer (*56*)) to measure mean intensity and cell apical area.

Side views from 40X image stacks along the posterior-anterior axis (Fig. 1D and 2D) were generated by a single line re-slice of the hyper-stacks across the dorsal midline. The side views were then smoothened with a Gaussian blur of 2 pixels (0.15μm) for representation.

For 100X image stacks, maximum intensity projections were used for measurements of fluorescence intensity and cell segmentation. Before projection, z-stacks were rotated, cropped and smoothened with a mean filter of kernel 0.5 pixels (0.04 μm) to increase the signal-to-noise (SNR). The 2D-projected stacks were then segmented and cells were tracked as previously described (*53*) using Ilastik (v.1.4) and Tissue Analyzer (*56*).

Side views from 100X (Fig. S7J) image stacks were generated by re-slicing a rectangle (6μm wide) across the dorsal midline on images smoothened with a mean blur of 1 pixel (0.08μm).

The distance of E-cad from the vitelline membrane (Fig. 1L, 2K and Fig. S7K) was measured on height map images carrying the information of the plane where E-cad junctions were most in focus in the stack. These height map images were generated from E-cad hyper-stacks using the Stack Focuser plugin with a kernel of 11 pixels (0.88μm). Height map images of E-cad were generated for all time points and a mask based on E-cad intensity in 2D max projection was applied to mask out regions where E-cad junctions were not present.

Kymographs for E-cad::GFP (Fig. 1M) along the posterior-anterior axis were obtained with a method similar to the generation of side views but using maximum intensity projection time series as an input for re-slicing. Kymographs are obtained using a single line of line width of 20 pixels (1.6μm) across the medio-apical region of the cell.

### Data analysis

#### Measurement of posterior endoderm displacement from brightfield imaging (Fig. S1C, Fig. S2C,E,G, *Fig. 4C*)

The extent of posterior endoderm displacement during gastrulation was measured from brightfield movies for 60 min after the time when the cellularization front reaches the basal side of the nuclei on the dorsal side of the embryo (Time 0). The cumulated displacement was measured by manually tracking the posterior edge of the invagination and by calculating the cumulative travelled distance of this point, l=√(x−x_0_)^2^+(y−y_0_)^2^, where (x_0_,y_0_) is its position at time 0. This was normalized to the maximum embryo length (L) measured as the distance between the anterior-most and posterior-most points of the embryo. Normalized posterior endoderm (PE) displacement = l/L.

#### Measurements of fluorescence intensity and cell shape

Measurements of fluorescence intensity and cell shape parameters were extracted using ImageJ and further analyzed with Matlab (R2022b including curve fitting toolbox, image processing toolbox, statistics and machine learning toolbox). For all intensity measurements, local background intensities were subtracted as previously described (*6*).

##### Measurement of the mean intensity of MyoII and apical area in 40X images (Fig. 1F, Fig. 2G)

Fluorescent intensity and apical surface area were measured on stack-focused projections of 40X image stacks. Individual cells were manually tracked by drawing ROIs (about 6 cells per embryo). The mean grey value and area were measured using ImageJ.

##### Measurement of the mean/integrated intensity of MyoII in 100X images (Fig. 1J,K, Fig. 2J, Fig. S7B)

Fluorescence intensities were measured on standard maximum intensity projections for 100X image stacks. For individual cell measurements, mean and integrated intensities were measured within a region of interest (ROI) obtained by automated segmentation and tracking and then shrunken by 10 pixels to remove junctional signals. Cell tracking was performed based on the E-cad signal in maximum intensity projections of E-cad. Time traces of MyoII and cell apical area in individual cells were registered based on the time when the apical area reached a maximum. This time is defined as time 0 in Fig. 1J,L, 2K and Fig. S7B.

#### Measurements of the invagination depth (Fig. 1G, Fig. 2F)

To measure the invagination depth a straight line was manually drawn in side views from the deepest point of the apical surface of the invagination to the overlying vitelline membrane. The distance measured at 8min (for Fig. 1G) or at 5min (for Fig. 2F) was plotted for the indicated conditions. Time 0 is defined by synchronizing embryos relative to the time of onset of cell divisions in the dorsal posterior (see embryo synchronization below)

#### Manual measurement of the number of cell rows activating MyoII during wave propagation (Fig. 1H, Fig.2H)

To measure the propagation of the MyoII wave, we manually recorded the time of MyoII activation in cells of the PZ along 3–4 rows of cells parallel the anterior-posterior axis in each embryo. For each embryo, a mean value was obtained by averaging the 3-4 rows and different embryos were averaged to obtain plots in Fig. 1H and 2H. Time 0 is defined as the time of MyoII activation in the first cell in the PZ for each embryo.

#### Space-intensity plot (Fig. 3B,C,E,F,H,I, Fig. S3C, Fig. 4C,E,J,K, Fig. S4F,G, Fig. 6B,F,I,J and Fig. S7D,E,F,H)

In Fig. S3C space intensity plots were measured from still images. First, a maximum intensity projection of 4 planes (2μm) was obtained. The Ptc signal was used as a proxy to define the posterior boundary of the *wg*Gal4 driven-Fog expression domain. The mean intensity of Fog along 3 lines with a thickness of 100 pixels (9 microns each) was measured from this posterior boundary (0 μm) and plotted as a function of distance along the A-P axis. The plots obtained were normalized (using min-max normalization) for representation.

In Fig. 4C space intensity plots were measured from still images. The mean intensity of Fog and MyoII was measured on maximum intensity projections of 4 planes (2μm) and an average background (across 3 embryos) was subtracted. The resultant mean intensity was plotted across the distance from the invagination along the A-P axis. To control that the observed profiles do not depend on a confocal sectioning effect, image stacks with the same z-settings were acquired using the 405nm laser line to capture embryo autofluorescence (grey background profile). Intensity profiles of the autofluorescence were measured to ensure that this value was constant. For Fig. 4E, the mean intensity of GFP and Fog was plotted similarly to Fig. 4C, without background subtraction.

In Fig. 3B,C,E,H, Fig. 6B,F,I,J and Fig. S7D,F space intensity plots were measured from registered time series where the position of the invagination front was kept constant. For this, the position of the front of the invagination was tracked manually over time using the ImageJ plugin (Manual tracking, https://imagej.nih.gov/ij/plugins/track/track.html). Each individual time point was then translated to keep the invagination front always in the same position. Then, an average intensity projection over time of the registered time series of MyoII was performed after background intensity subtraction. Additionally, the junctional MyoII signal was removed using a mask based on E-cad intensity above a threshold before the time average projection. The mean intensity of MyoII was then measured using 3-line profiles (200-pixel, 16 μm wide each) along the AP-axis of the embryo on the dorsal side. For each embryo, data were averaged across the 3-line profiles and plotted against the distance from the front of the invagination along the AP-axis. Data from all embryos were then pooled and normalized in GraphPad Prism (10.0.2/171) using min-max normalization. The value R^2^ in Fig. 3B,C quantifies the goodness of fit in GraphPad Prism. R^2^ (a fraction between 0.0 and 1.0) compares the fit of the one-phase exponential decay model to the fit of a horizontal line through the mean of all values on the Y-axis. R^2^ is computed as a ratio of sum of the squares of distances of the points from the best-fit curve determined by the fitting of the one-phase exponential decay model (SSresiduals) normalized to the sum of squares of the distances of the points from a horizontal line through the mean of all Y values (SStotal). R^2^ is calculated using the equation: R^2^= 1.0 – (SSresiduals/SStotal). R^2^ value closer to 1.0 indicates that the best-fit curve fits the experimental data.

For quantification of the length-scale (Fig. 3F,I, Fig. S7E), the averaged mean intensity profiles from individual embryos were fit with a one-phase decay exponential Y=(Y_0_ - a)*e^(-x/*λ*)^ + a, where Y0 is the measured intensity value at X=0, a is the plateau intensity value at X=∞ and λ is the decay length.

In Fig. 4K, Fig. S4G, Fig. 6F (dextran) and Fig. S7F, h space-intensity profiles were also measured from registered time series relative to the position of the invaginating front. Here, average projections over time of maximum intensity projections were obtained without local background subtraction. Average time projections were then used to measure the intensity using 3-line profiles of 100-pixel (9 μm) line width along the AP-axis of the embryo at the dorsal side.

In Fig. 4J and Fig. S4F, ratios of each line profile from one condition to all line profiles of the other condition were calculated. The average of all measured ratios was plotted against the distance from the front of the invagination.

In Fig. 6F and Fig. S7D,H the normalization is a min-max normalization.

#### Amplitude and range of MyoII and βPS (Fig. 3E,H, Fig. 6C,D,G,K-N and Fig. S7I)

For the amplitude, descriptive statistics of the space-intensity profiles of each embryo were obtained in GraphPad Prism and the maximum values for each embryo were plotted in Fig. 6B,I,J.

For the data in Fig. 6 and Fig. S7 the range was estimated as follows. The mean intensity and standard deviation (SD) of MyoII and β*PS* were measured in the region of very low/no activation in the embryo anterior of control embryos (the region from 50 to 60 μm anterior to the invagination front). An average of the values corresponding to 3 SDs above the mean for each control embryo was used as a global threshold of intensity to define activation for all conditions. The corresponding x-axis values were plotted as the range of activation for each embryo. For Fig. 6C and 6G, each embryo intensity profile was smoothened (80-90 neighbours on each size to average and 0^th^ order of the smoothing polynomial) in GraphPad Prism to achieve a precise value of the range. For Fig. S7I, a threshold of 10% of the normalized intensity was used to define a region of contact with the vitelline membrane.

For the data in Fig. 3 (Fig. 3E and 3H) the activation threshold was defined as the mean + 3 SDs estimated from the average curves of the controls in the region from 35 to 40 μm anterior to the invagination front, since the measurements did not cover a larger region. This global threshold (indicated by the dotted lines in Fig. 3E and 3H) was used to estimate the range in individual embryos for all conditions as above. Since the measurements did not cover regions beyond 40 μm anterior to the invagination front, in the text we indicated the lower limit of the possible range of MyoII activation for the mutant conditions.

#### Embryo synchronisation

Synchronization was performed to compare mutant to wild-type embryos and to register them in developmental time before data pooling and averaging.

For bright field movies (Fig. S1 and S2 for Fig. S4B-C), time 0 is defined as the time when the cellularization front reaches the basal side of the nuclei on the dorsal side of the embryos.

For 40X time lapses, synchronization is based on the time of onset of cell divisions in the dorsal posterior ectoderm. This phenomenon is independent of the morphogenesis of the posterior endoderm. Time 0 is defined as 22 min (for Fig. 1D,F) and 19 min (for Fig. 2G,H) prior to the first cell division in an embryo to coincide with the onset of MyoII activation in the primordium for control embryos.

For fluorescence experiments in fixed samples, embryos were staged manually based on similarity with events observed in time-lapse experiments.

### Fluorescence Correlation Spectroscopy (FCS)

#### Data acquisition

FCS measurements were performed at room temperature (22 °C) on an inverted Zeiss LSM880 system (Carl Zeiss, Oberkochen, Germany) using a Plan-Apochromat 100x, 1.4NA DIC M27 oil immersion objective. Fluorescence was excited with the 488 nm (Smog::GFP, VSV-G::GFP) or 514 nm (Fog::YFP, sec::GFP) line of an Argon laser in spot mode. Fluorescence was detected in the range of 491-700 and 516-700 nm (for measurements in Fig. 4G,H and Fig. S5A-E) or 491-558 and 516-558 nm nm (for measurements performed in Fig. 5I,J,L-O and Fig. S6A-D, and S6F-M) on a 32 channel GaAsP array detector operating in photon counting mode, with a pinhole set to one airy unit. Prior to FCS measurements and at embryonic stages preceding wave propagation, the vitelline membrane was pre-bleached to reduce autofluorescence background in the measurements. For this, the apical-most plane of the embryo (containing the vitelline membrane) was scanned in full field-of-view and continuous scan mode (scan speed 6) for 4 min with the 488 nm line at a power of ca. 15 µW or for 1.5 min with the 514 nm line at ca. 5 µW. Laser powers were measured at the exit of the objective. 4-5 FCS measurements per embryo were then acquired within the extracellular space of the invaginating furrow at an approximate depth of 5-10 µm below the vitelline membrane (spot I) during wave propagation. In addition, up to 10 measurements per embryo were performed at the apical surface of cells in the propagation zone (spot PZ), in a focal plane approximately 0.1 μm below the vitelline membrane. Each FCS trace was acquired for 30-60 s with a time resolution of 1.23 µs, at 0.2-2 µW excitation laser power for Fog::YFP and 0.2-6 µW for sec::GFP and Smog::GFP/ VSVG::GFP. This power range was selected to evaluate the potential bias of the estimated diffusion dynamics by light-induced photo-physical transitions of the fluorophores (see below) (*57, 58*). For oligomerization measurements of Smog::GFP and VSV-G::GFP, additional measurements of the residual autofluorescence background were performed in yw embryos, i.e. embryos not expressing any fluorescent construct, on the same day.

#### Data analysis

FCS measurements were manually exported as TIFF files using the ZEN software (Zen Black 2.3 SP1 FP3), imported and analyzed using custom-written MATLAB (The MathWorks, Natick, MA, USA; version R2020a) procedures previously described (*42, 59, 60*). To correct for signal decrease due to photobleaching, a two-component exponential function was fitted to the fluorescence time series and a correction formula was applied as previously described (*61*). For the analysis of Smog and VSV-G in cells of the PZ, an additional correction was applied to remove residual background from the vitelline membrane. To this end, background intensity time traces acquired in multiple pre-bleached yw embryos were averaged and a two-component exponential fit was fitted to the average intensity time trace. The obtained fit function was then subtracted from the intensity time trace of each individual measurement acquired in the actual sample, i.e. embryos expressing Smog::GFP or VSV-G::GFP. Due to local variations in the background intensity, this correction procedure occasionally led to very low or even slightly negative intensity values, inducing high noise particularly in the brightness readout. We therefore removed all measurements with average intensities below 10 kHz after background subtraction (typically 10-20% of measurements at maximum). Generally, background correction by subtraction is valid in the case of non-correlated background, which we confirmed by FCS analysis of the background measurements (as also shown in (*37*)). Note that the applied correction scheme accounts, on average, for temporal decay of the background intensity due to photobleaching. In fact, intensity time traces of Smog::GFP or VSV-G::GFP did not show substantial photobleaching after background subtraction.

From the resulting intensity time trace F(t), the autocorrelation function (ACF) was calculated as follows, using a multiple tau algorithm:

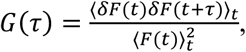

where *δF*(*t*) = *F*(*t*) − 〈*F*(*t*)〉*_t_*.

To avoid artefacts caused by rarely occurring long-term instabilities, ACFs were calculated segment-wise (5-10 segments) and then averaged. Segments showing clear distortions were manually removed from the analysis (*62*). In addition, the first segment was always removed since it was occasionally corrupted by residual intensity changes due to temporal variations of background photobleaching that were not properly captured by the average background correction.

For measurements inside the invagination (spot I), a model for three-dimensional diffusion and Gaussian confocal volume geometry was fitted to the average ACFs (*59*):

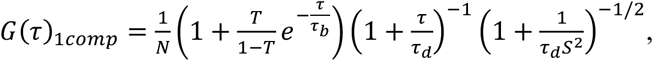

Here, *N* denotes the particle number, *τ_d_* the diffusion time, *T* is the fraction of fluorophores undergoing photophysical transitions with an average time constant *τ_b_* (see details below), and *S* is the structure parameter. From the diffusion time, the diffusion coefficient *D* was determined by 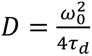. The waist *ω*_0_ was calibrated from FCS measurements of Alexa Fluor® 488 (Thermo Fisher Scientific, Waltham, MA, USA) dissolved in water at 20 nM, using the previously determined diffusion coefficient of 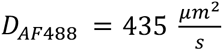 Ref. (*63*). Calibration measurements were performed at ca. 1 µW (at 488 nm) or 10 µW laser < power (to compensate for suboptimal excitation of AF488 at 514 nm) and 2 µm depth to minimize aberrations. The structure parameter was fixed to the average value determined in calibration measurements, typically around 6.

Under the assumption of an aqueous environment and globular protein conformation, the relative diffusion dynamics of two proteins (i.e. diffusion coefficients D_1_ and D_2_) of different size (i.e. of molecular weights MW_1_ and MW_2_) diffusing freely in 3D can be estimated using the Stokes-Einstein relation, 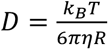 (with *k*_B_denoting the Boltzmann’s constant, T the temperature and *η* the viscosity of the solvent, and R the proteins hydrodynamic radius) and the fact that the hydrodynamic radius approximately scales with the cubic root of the molecular weight (for proteins of similar mass density). Thus, 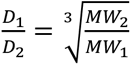. Using secreted GFP (molecular weight of ca. 27 kDa) as a ruler, it is thus expected that Fog::YFP (molecular weight of ca. 127 kDa) diffuses about 1.7 times slower than secreted GFP. With careful analysis of photophysical effects (see details below), a relative ratio of 1.6 was obtained with the determined diffusion coefficients of 55 µm^2^/s for Fog::YFP and 87 µm^2^/s for sec::GFP in the invagination.

For measurements in the propagation zone (spot PZ), a two-component diffusion fit model was applied:

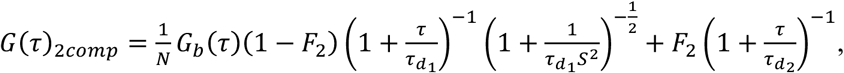

where *G_b_*(*τ*) denotes the photophysics term, 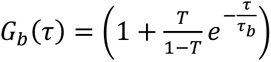, and *F_2_* the fraction of the second component. The first component is attributed to particles diffusing freely in 3D and the second component to diffusion in a 2D plane (*64*), corresponding to particles that diffuse extracellularly in the space between the vitelline membrane and the apical cell surface but potentially bind to receptors in the apical membrane resulting in (transient) 2D diffusion, a behavior that has often been observed for extracellularly diffusing morphogens (*60, 65, 66*). For measurements of membrane-bound proteins, i.e. Smog and VSV-G, the 2D diffusion term corresponds to molecular diffusion in the apical membrane, while the 3D diffusion term corresponds to intracellular background, as previously discussed (*37*).

To minimize the number of free fit parameters, a single photophysics term was included in the ACF fit models given above. However, the fluorescent protein tags, GFP and SYFP2, utilized in this study exhibit multiple (at least two) photophysical transitions (*57, 58, 62, 67*): 1) triplet state transitions occurring on the µs time scale, 2) (light-induced) flickering occurring on the 10-100 µs time scale. Thus, estimates of fast diffusion dynamics characterized by fluctuations on similar time scales might be biased. Indeed, fitting ACFs with maximal time resolution (fit routine 1) resulted in photophysics time constants of ca. 1 µs and illumination intensity-dependent diffusion coefficients (see Fig. 4G,H, Fig. S5B). To obtain unbiased diffusion coefficients, the values were interpolated to zero illumination power, according to a previous study (Vámosi et al. 2016). To cope with low signal-to-noise ratio of individual ACFs in this analysis, ACFs of several measurements acquired at the same power in multiple embryos were first averaged and the 3D diffusion model fitted to the average ACF (Fig. 4H). This was particularly crucial for sec::GFP, which is ca. 7-fold less efficiently excited as Fog::YFP at the same 514 nm laser power. A comparison of the diffusion coefficients obtained by fitting individual ACFs at comparable effective excitation (i.e., 0.25 µW for Fog::YFP vs. 2 µW for sec::GFP) is shown in Fig. 4G.

Measurements in the propagation zone (spot PZ) had to be performed at higher excitation power (i.e. 1-2 µW) to maximize signal-to-noise ratio and minimize acquisition time with respect to the movement of the propagating wave. To minimize bias in estimating the fast-diffusing component, an alternative fit routine was applied (fit routine 2, Fig. S5B), in which fitting was restricted to lag times larger than 5 µs. For measurements of Fog::YFP inside the invagination (spot I), the photophysics term then converged to time constants of 30-80 µs with a fraction of 8% at the lowest and 30% at the highest laser power. The resulting diffusion coefficients were almost independent of excitation power, showing that this fit model captures well the flickering contribution and hence minimizes the bias in the estimated diffusion dynamics (Fig. S5B). The second fit routine was therefore applied to analyze all measurements performed in the propagation zone (spot PZ). From the determined particle number *N*, the protein concentration *c* was estimated, 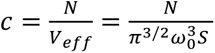, where 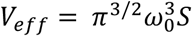 is the effective detection volume. It should be noted that for measurements of Fog diffusing in the propagation zone, this formula and the applied FCS fit model provide an approximation because the physical space in which molecules diffuse extracellularly is confined by the vitelline membrane at the top and the apical cell surface at the bottom.

For oligomerization measurements of the membrane proteins Smog and VSV-G, the two-component fit was dominated by the 2D diffusion term, corresponding to a fraction of 60-80% of molecules and diffusion times of ca. 5-100 ms. The 3D diffusion term converged to faster diffusion times of few hundred µs to few ms. The photophysics term converged to decay times of ca. 5-50 µs.

#### Molecular brightness measurements using FCS

The molecular brightness was quantified from the average fluorescence intensity and the particle number determined from the FCS analysis, 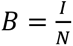. For a mixture of a monomeric species (brightness *B_1_*) and an oligomeric species (i.e. n-mers, brightness *B_n_=nB_1_*), the average brightness is calculated as follows, 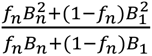, where *f_n_* is the relative fraction of the oligomeric species and *f_1_* = 1-*f_n_* the monomeric fraction(*42*). Inverting this equation allows computing the oligomeric fraction, 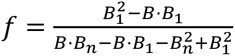. Previous studies have reported that fluorescent protein properties strongly affect brightness-based oligomerization measurements. In particular, dark fluorophore states have to be taken into account, which can effectively be modelled with a single parameter, the fluorescence probability *p_f_* (Dunsing et al. 2018). For an n-mer, the brightness *B_n_* in the equation for the oligomer fraction is then calculated as follows, 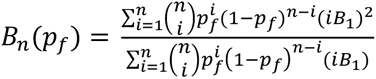, assuming that all fluorescent protein subunits are independent and in a fluorescent state with a probability *p_f_*.

In our analysis, we first calculated the brightness of monomeric GFP assuming that VSV-G::GFP is exclusively present as trimers and then used this value to calculate the oligomer fraction for a putative monomer-oligomer mixture of Smog::GFP, given the experimentally determined average brightness B. We calculated oligomer fractions for *p_f_* = 1 (i.e. absence of dark GFP states, since they have not been characterized in Drosophila embryos yet) and *p_f_* = 0.7 (i.e. the value that has been previously measured for GFP in mammalian cells (*42*)).

#### Gradient analysis using FCS

FCS measurements were performed at the apical cell surface at increasing distances from the moving invagination. Before each measurement, a confocal image capturing the position of the invagination front and the anterior part of the embryo was acquired where the position of the FCS spot was marked. In the analysis, the position of the invagination front was manually determined in the image. Then, FCS and image metadata were extracted (using custom-written MATLAB code) to determine the distance of the FCS spot from the invagination for each FCS measurement. Estimates for the relative fractions (F_fast_ and F_slow_) of fast and slow diffusing components and molecular brightness values resulting from FCS analysis were pooled for several measurements in each embryo and across multiple embryos. These values were then plotted as a function of distance from the invagination.

### Statistics

For all experiments, data points from different cells, measurements or embryos from at least two independent experiments were pooled to estimate the plotted mean, s.d. and s.e.m.

In order to determine whether the two datasets were significantly different, an unpaired *t-test* was performed using GraphPad Prism. It was assumed that the datasets are sampled from Gaussian populations but have unequal variances (Welch’s correction). No statistical method was used to predetermine the sample size. The experiments were not randomized, and the investigators were not blinded to allocation during experiments and outcome assessment.

#### Repeatability

All measurements were performed in 3-9 embryos. In experiments involving living embryos, we considered each embryo as an independent experiment. In immunostainings, independent experiments correspond to distinct staining procedures. Representative images, which are shown in Fig. 1-6 and Suppl. Fig. 1-7 were repeated at least twice and up to more than ten times.

#### Data availability

All the data supporting the findings of this study are available within the paper. Raw image data are available upon reasonable request.

---

**Table S1.**
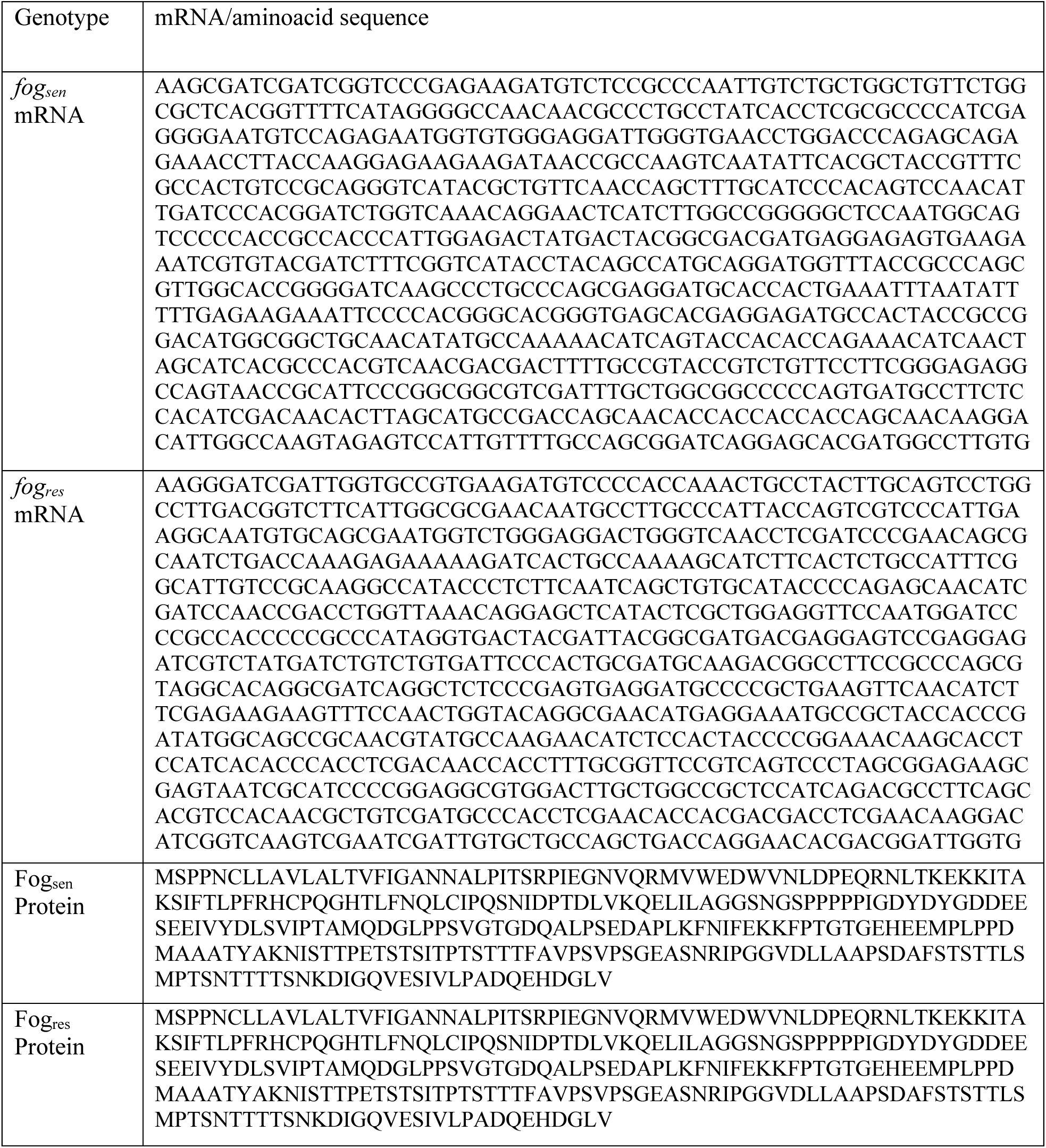
Alignment of the nucleotitidic (mRNA) and aminoacidic sequences of the original (RNAi-sensitive, *fog_sen_*) and the RNAi-resistant (*fog_res_*) versions of *fog*. Nucleotides were modified using codon redundancy across the entire *fog* dsRNA sequence, starting from −24bp before the ATG up to +834bp after the ATG (V279). The aminoacidic sequence remains the same, thus generating an RNAi-resistant version of *fog*.

## Notes

### Competing Interest Statement

The authors have declared no competing interest.

